# mTOR dysregulation induces IL6 and paracrine AT2 cell senescence impeding lung repair in lymphangioleiomyomatosis

**DOI:** 10.1101/2024.05.03.592025

**Authors:** Roya Babaei-Jadidi, Debbie Clements, Yixin Wu, Ken Chen, Suzanne Miller, Manuela Plate, Kyungtae Lim, Rachel Chambers, Emma L Rawlins, Yan Xu, Simon R Johnson

## Abstract

Lymphangioleiomyomatosis (LAM) is a rare disease which causes lung cysts and respiratory failure. TSC2 deficient ‘LAM cells’ with dysregulated mTOR signalling form nodules with fibroblasts causing lung injury. We examined if mTOR dysregulation could induce senescence and impair responses to lung injury. Senescence markers p21 and p16 were increased in LAM lungs and co-localised with alveolar type 2 cells. The SenMayo senescence gene panel was upregulated in LAM alveolar type 2 cells with senescence supressed by mTOR inhibition in patients. LAM cell / fibroblast spheroid cultures induced senescence markers in alveolar type 2 cell organoids, altered their growth and delayed epithelial scratch wound repair. Upstream regulator analysis predicted alveolar type 2 cell IL6 receptor activation. IL6 was produced by LAM cells, induced p16 and p21 in alveolar type 2 cells, inhibited epithelial wound resolution and was overexpressed in LAM patient serum where it was related to lung function. Wound repair in the presence of TSC2 null LAM cell / fibroblast spheroids was enhanced by the IL6 receptor antagonist Tocilizumab. Our findings show TSC2 loss induces senescence and IL6 production which are associated with impaired lung repair. Targeting IL6 signalling in parallel with mTOR inhibition, may reduce lung damage in LAM.

## Introduction

Lymphangioleiomyomatosis (LAM) is a rare multisystem disease that almost exclusively affects women. Most symptoms are caused by the progressive accumulation of lung cysts which lead to pneumothorax, dyspnoea and respiratory failure. The disease may also be associated with lymphatic obstruction and a propensity to develop the benign mesenchymal tumour, angiomyolipoma^1^. LAM can occur both sporadically and also in women with the genetic disease tuberous sclerosis complex (TSC) where LAM is a major cause of death and disability in adults^2^. The key pathological feature of the disease is the LAM cell, a clonal population of cells with loss of function of one of the TSC genes, most commonly *TSC2*. TSC gene loss results in dysregulated mechanistic target of rapamycin (mTOR) signalling leading to LAM cell expansion, migration, protection from cell death and altered metabolism^1^. LAM cells in the lung parenchyma proliferate locally and attract wild-type cells to form ‘LAM nodules’, clusters of cells comprising LAM associated fibroblasts (LAF), lymphatic endothelial cells, T cells and mast cells analogous to a tumour stroma^3–6^. As the disease progresses, LAM nodules become more complex increasingly recruiting wild-type cells which is associated with falling lung function^7^.

One of the striking features of LAM is the unusual cystic destruction of the lung parenchyma, with lung cysts present even when LAM cells are sparse^7^. The mechanism of lung damage is unknown although LAM nodules express matrix degrading proteases including cathepsin K^8^, matrix metalloproteinases^9^ and the plasmin system^10^ which are thought to cause lung cyst formation. In addition, knockdown of *TSC2* in mesenchyme also results in alveolar enlargement in mice which is dependent on Wnt pathway activation^11^. Lung repair after injury is characterised by the expansion of alveolar type 2 (AT2) cells, the resident alveolar stem cell, which differentiate to replace damaged alveolar type 1 (AT1) cells to repair the gas exchange surface of alveoli. AT2 cell stemness is maintained by Wnt signalling from adjacent PDGFRα-Axin2 expressing fibroblasts^12^ ^13^. Loss of this signal by alveolar injury induces a repair program causing differentiation of AT2 into AT1 cells^14^ ^15^. Single cell RNA sequencing (scRNAseq) of human lung injury and organoid models of the repair process have revealed cells with both AT1 and AT2 markers. These intermediate cells, termed ‘pre-alveolar type-1 transitional cell state’ (PATS), have also been identified in single cell analyses and dual label immunohistochemistry in LAM^16^. PATS are unable to differentiate into AT1 cells and may contribute to impaired repair in lung disease^17^. Their presence in LAM lung suggests that in addition to proteolytic lung injury there may be a co-existent failure of lung repair.

Senescence is a state of irreversible cell cycle arrest, which in health, prevents the replication of damaged cells, suppressing tumorigenesis and limiting scar formation. Senescence occurs due to repeated cell division resulting in telomere shortening (replicative senescence) or by DNA damage resulting from reactive oxygen species (ROS) or other stressors including cigarette smoke^18^. Senescent cells tend to be large with flat morphology and increased lysosomal senescence-associated beta-galactosidase (SAβgal) activity. DNA damage, sensed by activation of the DNA repair kinase ATM, activates p53 and p21 to suppress the cell cycle. Other stressors can directly activate another cyclin inhibitor, p16^INK4a^ to cause cell cycle arrest^19^. Importantly, senescent cells remain metabolically active and secrete pro-inflammatory cytokines, chemokines and proteases. This senescence-associated secretory phenotype (SASP) acts in a paracrine manner to induce senescence in neighboring cells^20^ ^21^. The accumulation of senescent cells in disease and aging, and the pro-inflammatory SASP result in reduced organ repair, altered immune function and accelerated tissue damage in response to injurious stimuli. Suppression of senescence or the SASP, including by inhibition of mTORC1, activation of adenosine monophosphate kinase (AMPK), activation of sirtuins or targeted removal of senescent cells by senolytic therapies all extend lifespan in model organisms^22^. Activation of mTORC1, either by reactive oxygen species, AMPK inhibition or genetic loss of PTEN or TSC1/2, is central to senescence; downregulating sirtuins 1 and 6 to both suppress DNA repair and induce the SASP via NFκB signaling^23^. We therefore hypothesised that mTOR dysregulation in LAM cells might induce senescence of other LAM nodule components and via the SASP induce senescence in surrounding AT2 cells impairing lung repair processes.

## Results

### Markers of senescence are present in LAM lungs

To determine the presence of senescent cells in LAM lung tissue, we initially examined gene expression of two canonical senescence markers, the cyclin dependent kinase (CDK) inhibitors *CDKN1A* (p21) and *CDKN2A* (p16), using the LAM cell atlas, an open-source data interface comprising single cell RNA sequencing data from 13 lung samples^24^. p21 was ubiquitously expressed throughout the LAM lung including in *TSC* null LAM^core^ cells. Within the alveolar epithelial cell population, p21 is most enriched in AT1/AT2 transitional (PATS) cells. p16 expression was lower than p21 across all cell types, although >10% of LAM^CORE^ cells were p21 positive, representing a 2-fold enrichment compared with other cell types other than macrophages (Figure 1A).

**Figure 1.**
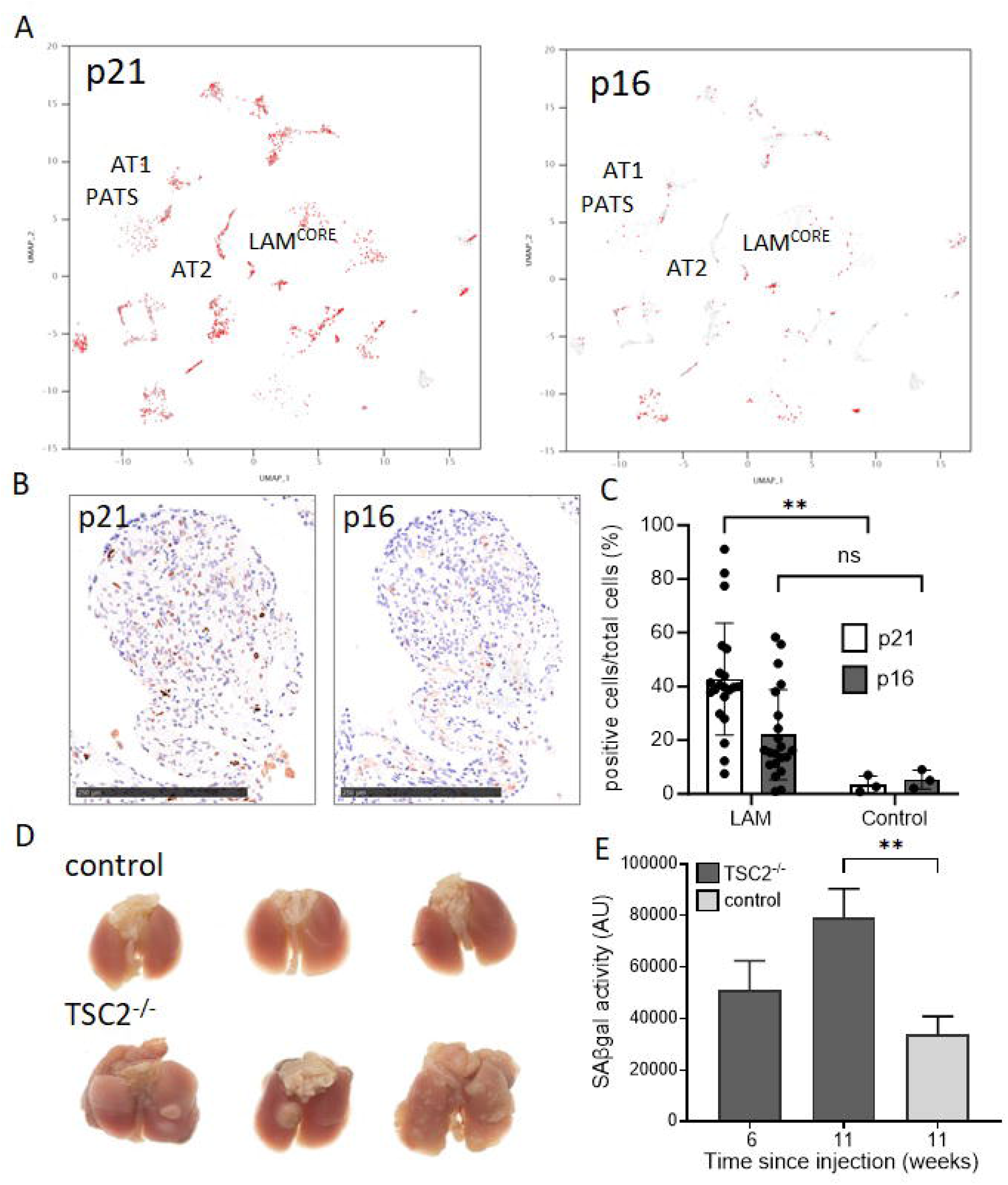
Markers of senescence are present in LAM lung. (A) Single cell RNA sequencing of LAM lung tissues showing expression of p16 and p21 in LAM lung populations using the LAM cell atlas^24^. Left panels show expression of positive cells for p21 and p16 in each cluster, right panels show the fold change and significance for LAM compared with control lung. Lower panel shows cell types present in LAM lung samples. AT1 = alveolar type 1 cells, AT2 = alveolar type 2 cells, PATS = pre-alveolar type-1 transitional cell state. (B) Immunostaining of LAM nodules for p16 and p21 protein, representative images of 21 lung sections. (C) Quantification of p16 and p21 expressing cells from 21 LAM and three healthy control lung tissues, using QuPath software^48^. (D) Gross appearance of albino black 6 mouse lungs 11 weeks after TSC2^−/−^ cell injection or sham injection (control). Tumour nodules are visible in TSC2^−/−^ cell treated lungs. (E) Senescence associated beta galactosidase (SAβgal) activity in albino black 6 mouse lungs after TSC2^−/−^ cell injection or control. Graph shows mean (SD) of three animals per group. **p<0.01.

As p21 and p16 proteins are mostly regulated by ubiquitin mediated degradation rather than transcription, we used immunohistochemistry to examine p16 and p21 protein in 21 LAM lung and 3 control lung samples. p21 and p16 were expressed in 42% and 20% of cells within LAM lungs respectively with p21 significantly more abundant in LAM than control lung tissues (Figure 1B, C and supplementary figure 1). We next examined the activity of senescence associated beta galactosidase (SAβgal) in a *Tsc2* null murine homograft model of LAM. Immuno-edited *Tsc2* null murine renal tumour cells were injected into the tail veins of immunocompetent black 6 mice with control animals undergoing sham injections with tissues harvested at six and 11 weeks post cell injection. Tumour nodules were visible in *Tsc2* null homografts and SAβgal activity in lung lysates was greater than two-fold higher than control animals after 11 weeks (Figure 1D, E).

### LAM nodules contain senescent cells

To characterise the senescent cell types in the LAM nodule stroma we performed dual immunostaining for p21 and p16 with the LAM cell markers PNL2 and GP100 respectively. Control lungs did not stain for either LAM cell marker (Figure 2A and supplementary figure 2). p21 was significantly increased in LAM nodules compared with control lungs, p16 tended to be more common in LAM but not significantly so. Only occasional cells co-expressed p21 or p16 proteins and the LAM cell markers, suggesting the majority of senescent cells within nodules were non-LAM cell types (Figure 2B). Examining lung function at the time of tissue sampling showed that p21 and p16 expressing cells, were present at similar levels in patients of all degrees of disease severity (Supplementary figure 3).

**Figure 2.**
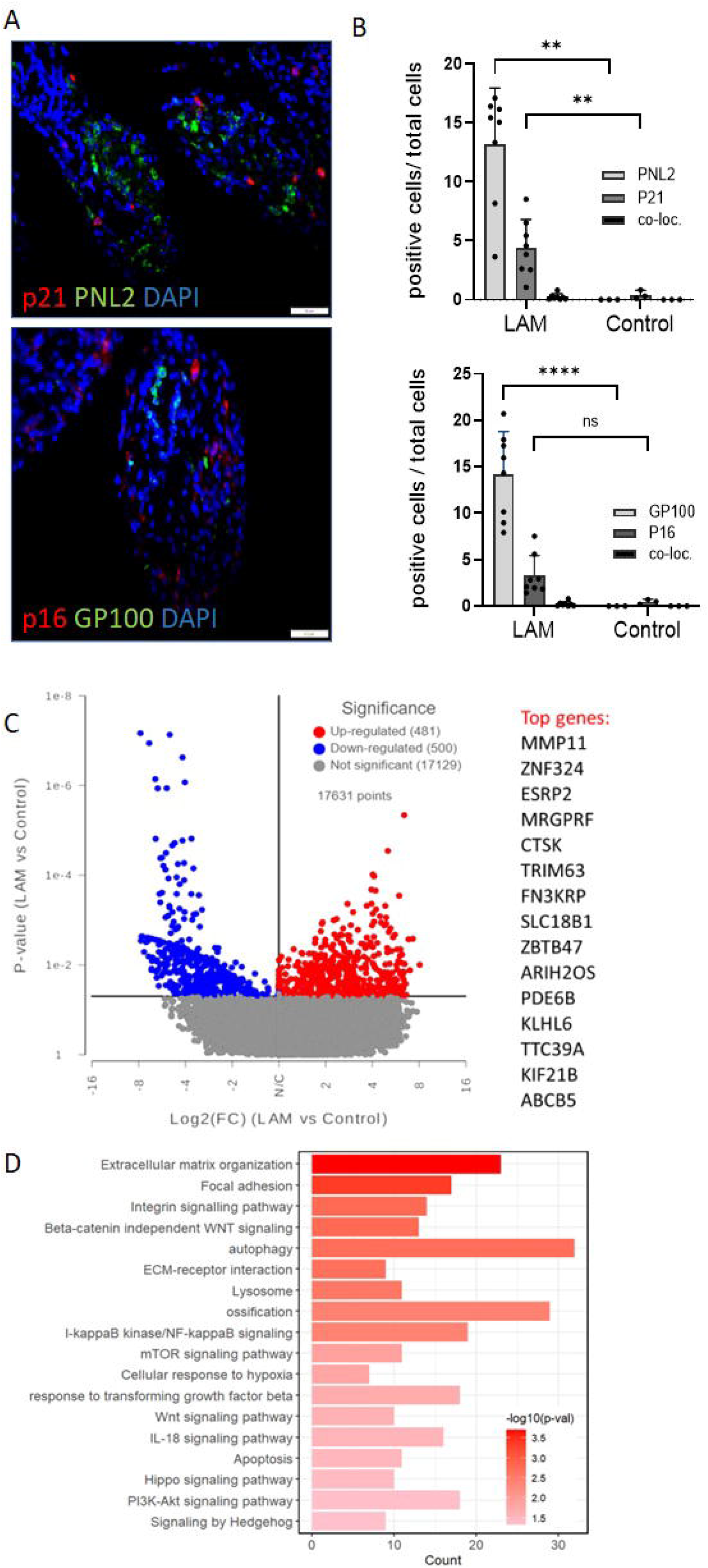
Senescence in LAM nodules. (A) Representative images of dual label immunohistochemical staining of p16 and p21 and the LAM cell markers PNL2 and GP100 respectively. (B) Quantification of senescence and LAM cell markers in eight LAM and three control lungs. Graphs show the mean (SD) percentage of positive, compared with all cells in the region of interest and the percentage of cells expressing both markers (co-loc). (C) Volcano plot showing differentially expressed genes assessed by RNA sequencing in LAM nodules isolated by laser capture micro-dissection from 19 women with LAM compared with healthy control lungs. The 15 most significantly increased genes are listed. (D) Pathway analysis of laser captured LAM nodules showing the most strongly increased pathways. **p<0.01, ****p<0.0001.

To understand the processes driving senescence we used laser capture micro-dissection to isolate LAM nodules from lung tissue from 19 women with LAM (Table 1) and analysed gene expression by RNA sequencing. 481 individual genes were upregulated in LAM nodules and 500 downregulated compared with healthy lung tissue (Figure 2C and supplementary table 1). Pathway analysis showed alterations in processes known to underlie LAM biology including mTOR signalling, autophagy and hypoxia. In addition, there was a strong extracellular matrix signal including ECM organisation, ECM receptor interaction, TGFβ response and integrin signalling. Genes associated with apoptosis and IL-18 signalling were also increased (Figure 2D).

**Table 1.** Clinical features of patient samples for laser capture microdissection.

| Patient characteristics |  |  |  | Disease at time of tissue sample |  |  | Lung function loss |  | Disease features |  |  |  |  |
| --- | --- | --- | --- | --- | --- | --- | --- | --- | --- | --- | --- | --- | --- |
| Age (yrs) | Race | Sporadic or TSC-LAM | Menopause | FEV<sub>1</sub> | TL<sub>CO</sub> | Disease duration (months) | ΔFEV<sub>1</sub> | ΔTL<sub>CO</sub> | Presentation | Ever had pneumothorax | Angiomyolipoma present | Lymphatic involvement | VEGFD (pmol/ml) |
| 49 | Caucasian | sporadic | yes | 18.7 | 33.4 | 228 | -0.105 | -0.370 | angiomyolipoma | yes | yes | no | 666 |
| 49 | Caucasian | sporadic | no | 77.5 | 60.8 | 240 | NA | NA | pneumothorax | yes | no | no | 2323 |
| 20 | African | TSC-LAM | no | 50.4 | 56.0 | 3 | -0.075 | -0.163 | pneumothorax | yes | yes | no | 289 |
| 30 | Caucasian | TSC-LAM | no | 76.0 | 57.1 | 1 | -0.092 | -0.132 | dyspnoea | yes | yes | no | 1028 |
| 69 | Caucasian | sporadic | yes | 83.1 | 45.3 | 228 | NA | NA | pneumothorax | yes | no | no | 397 |
| 44 | Caucasian | sporadic | yes | 49.5 | 40.0 | 72 | -0.029 | -0.082 | pneumothorax | yes | yes | no | 1326 |
| 31 | Caucasian | sporadic | no | 94.9 | 67.3 | 12 | -0.142 | -0.247 | angiomyolipoma | yes | yes | no | 702 |
| 28 | Caucasian | sporadic | no | 78.4 | 56.8 | 1 | NA | NA | other respiratory | no | no | yes | 4287 |
| 49 | Caucasian | TSC-LAM | no | 69.8 | 68.8 | 60 | -0.166 | -0.275 | chance finding | yes | yes | no | NA |
| 37 | Iranian | sporadic | no | 66.3 | 72.5 | 1 | -0.063 | -0.261 | pneumothorax | yes | yes | yes | 1163 |
| 48 | Caucasian | sporadic | yes | 78.0 | 73.0 | 12 | NA | NA | pneumothorax | yes | no | no | 3765 |
| 21 | Caucasian | TSC-LAM | no | 35.2 | 37.9 | 12 | -0.089 | -0.656 | angiomyolipoma | yes | yes | no | 3590 |
| 40 | Caucasian | sporadic | no | 89.6 | 74.9 | 35 | -0.258 | -0.731 | pneumothorax | yes | no | no | 586 |
| 36 | Caucasian | sporadic | no | 76.0 | 54.7 | 72 | -0.014 | -0.133 | pneumothorax | yes | no | no | 645 |
| 30 | West Indian | sporadic | no | 67.9 | 37.3 | 48 | NA | NA | dyspnoea | no | no | no | 967 |
| 41 | Caucasian | sporadic | no | 87.2 | 81.8 | 6 | -0.028 | -0.094 | pneumothorax | yes | no | no | 416 |
| 30 | West Indian | sporadic | no | 67.9 | 37.3 | 48 | NA | NA | dyspnoea | no | no | no | 967 |
| 21 | Caucasian | TSC-LAM | no | 35.2 | 37.9 | 12 | -0.089 | -0.656 | angiomyolipoma | yes | yes | no | 3590 |
| 55 | Caucasian | sporadic | yes | 91.5 | 90.3 | 23 | -0.059 | -0.155 | dyspnoea | yes | no | no | 834 |

FEV<sub>1</sub> / TL<sub>CO</sub> = percent predicted value for forced expiratory volume in 1 second / transfer of lung carbon monoxide at the time of tissue biopsy. ΔFEV<sub>1</sub> = rate of loss of FEV<sub>1</sub> (litres/year). ΔTL<sub>CO</sub> = rate of loss of TL<sub>CO</sub> (mmol/min/kPa/year). VEGF-D = serum vascular endothelial growth factor type D. NA = not available

### AT2 cells are senescent in LAM

We hypothesised that senescent cells within LAM nodules would produce a SASP response and induce senescence in neighbouring cells including the AT2 population. In LAM, AT2 cells are observed surrounding LAM nodules and lung cysts in addition to their normal alveolar niche^16^ ^25^. To quantitate senescent AT2 cells we performed dual immunohistochemical staining in 15 LAM and five control lung tissues using the AT2 cell marker pro-surfactant protein C (pro-SPC) and either p21 or p16. Healthy control lung AT2 cells did not express either p16 or p21, whereas populations of both nodule-associated and parenchymal AT2 cells expressed p16 and p21 proteins in LAM lungs (Figure 3A, B and supplementary figure 2).

**Figure 3.**
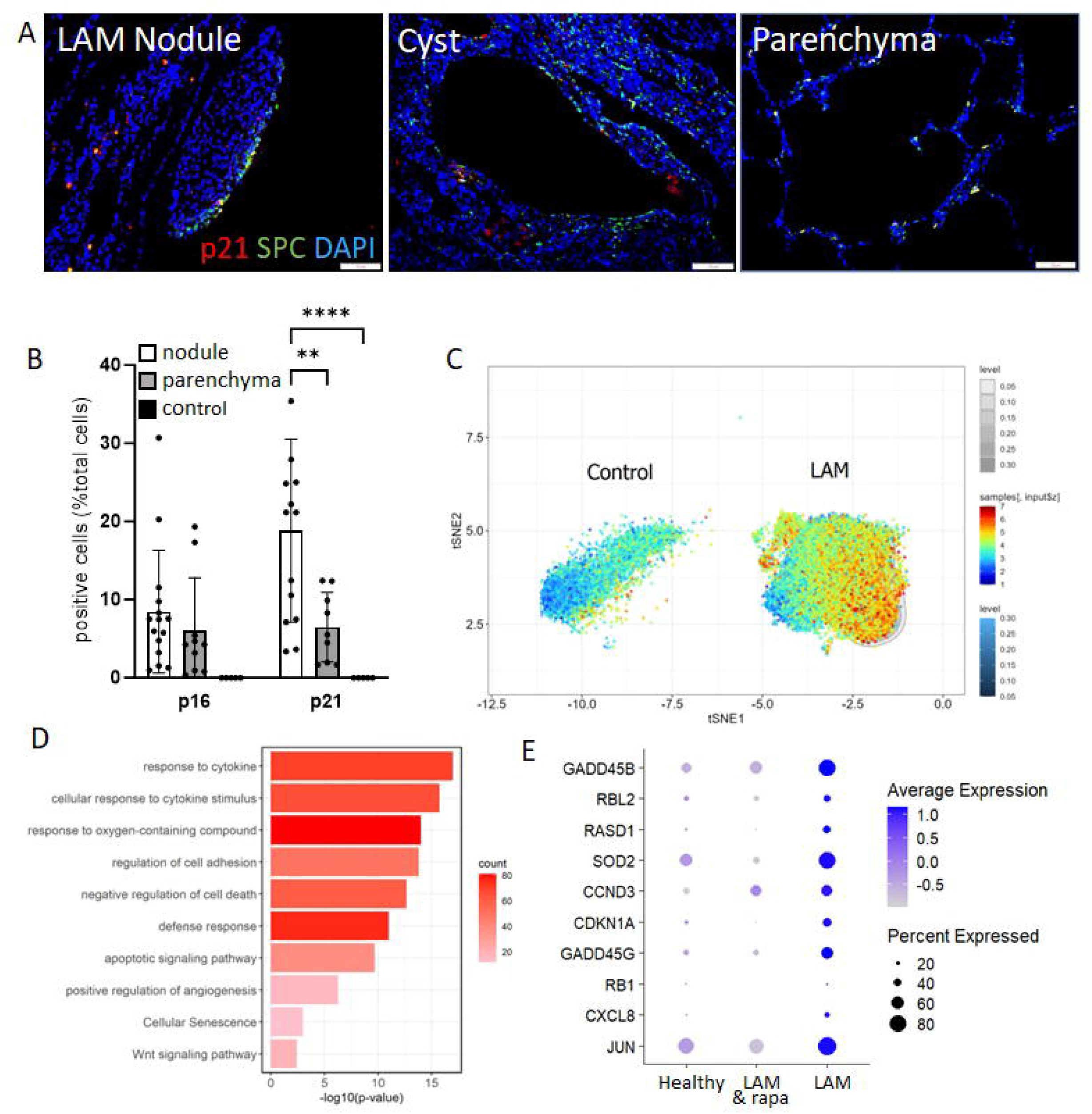
AT2 cells are senescent in LAM. (A) Representative images of dual label immunohistochemical staining of p21 and the AT cell marker surfactant protein C (SPC) around LAM nodules, in the walls of lung cysts and more normal areas of lung parenchyma. Similar findings were observed with p16 (not shown). (B) Quantification of co-expression of SPC and senescence markers in 11 LAM and five control lungs. Graphs show the mean (SD) percentage of positive, compared with all cells in the region of interest around LAM nodules in the lung parenchyma and in healthy control lungs. (C) Expression of the Senmayo geneset in control and LAM AT2 cells in the LAM cell atlas^24^. (D) Differentially expressed pathways in LAM compared with healthy control AT2 cells. (E) Expression of a panel of senescence associated genes from single cell RNA sequencing from LAM and healthy control AT2 cells from the LAM cell atlas. Single cell RNA sequencing from a single lung from a patient with LAM treated with the mTOR inhibitor rapamycin (LAM & rapa) showing suppression of senescence associated genes. **p<0.01, ****p<0.0001.

As senescence is a complex cellular response, which cannot be robustly represented by a single marker we examined the SenMayo senescence gene signature, a validated panel of 125 genes responsive to senescence in a range settings^26^. The SenMayo gene panel was significantly enhanced in LAM derived AT2 cells when compared with normal lung AT2 cells (Figure 3C and supplementary table 2). Examining pathways associated with the differentially expressed genes (DEGs) in LAM AT2 cells, we observed evidence of cytokine stimulation consistent with SASP activity, senescence, apoptosis and Wnt signalling (Figure 3D). To determine the effect of mTORC1 suppression on AT2 cell senescence in human LAM we compared a panel of AT2 cell senescence associated genes in control lungs, rapamycin naive women with LAM and a LAM patient treated with rapamycin. AT2 cell senescence associated genes were increased in LAM and were almost completely normalised in the patient treated with rapamycin (Figure 3E).

### SASP mediated senescence induction is mTOR dependent

To study the interactions between LAM nodule components and the alveolar epithelium we established a series of *in vitro* co-cultures incorporating co-cultures of LAM patient derived *TSC2* null 621-101 or control *TSC2* add-back 621-103 cells with primary LAFs isolated from LAM lung tissue. We initially characterised the senescence response in low serum co-cultures after 14 days and observed LAF / 621 cell interactions induced a senescent phenotype characterised by enlarged flattened cells with increased numbers of lysosomes, increased SAβgal activity, p16, p21 protein expression and nuclear translocation of p16 (Figure 4A and supplementary figure 4A, B). Using SAβgal as a reporter of the senescent phenotype we showed that both LAM cells and LAFs could mutually induce senescence in the other cell type. Senescence induction was TSC2 dependent with *TSC2* null 621-101 cells having higher levels of SAβgal than *TSC2* addback 621-103 cells and senescence induction by LAFs in 621 cells also being *TSC2* dependent with only the *TSC2* null 621-101 cells able to increase SAβgal expression in LAFs (Figure 4A). Bulk RNA sequencing of the LAM cells from co-cultures showed an induction of senescence initiating genes including *p16, p21, TP53, RBL1, RBL2* in a *TSC2* dependent manner (Supplementary figure 4C).

**Figure 4.**
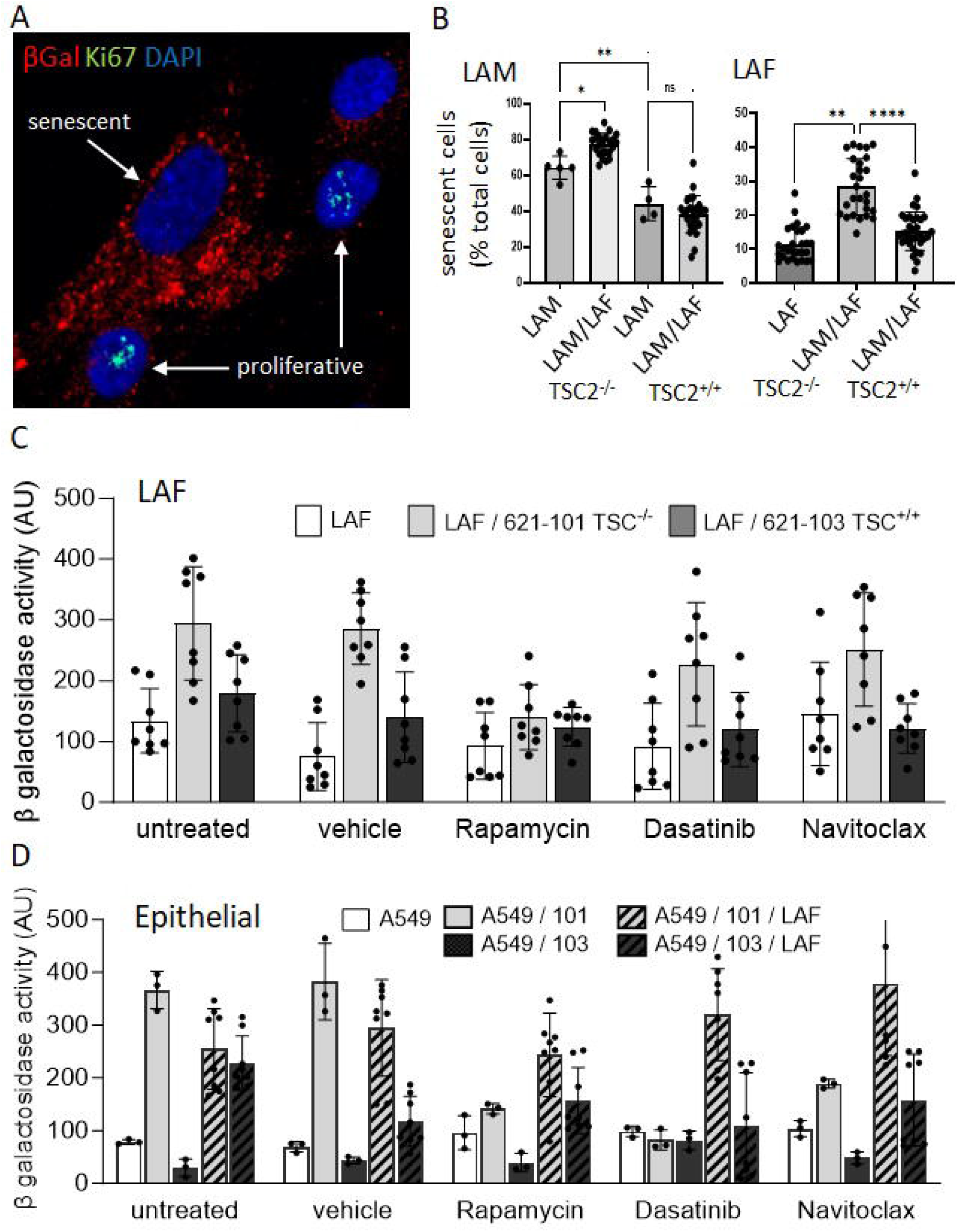
Cell-cell interactions induce senescence dependent on mTOR dysregulation. (A) Immunofluorescent staining of 621-101 cells incubated with LAFs for 14 days. Senescent cell highlighted is enlarged with increased senescence-associated beta-galactosidase (βgal) positive lysosomes when compared with proliferative cells with nuclear ki67 staining. (B) Interactions between LAM derived 621 cells (LAM), LAM associated fibroblasts (LAF) and LAM/LAF co-cultures assayed after 14 days. Graphs show mean (SD) percentage of LAM (left panel) or LAF (right panel) when interacted in culture cells expressing senescence associated beta galactosidase. TSC2^+/+^ are control, addback 621 cells used to show the effect of mTOR dysregulation on the induction of senescence. (C) Mean (SD) senescence associated beta galactosidase activity in LAFs when cultured with 621 cells and the effect of rapamycin and the senolytic drugs Dasatinb and Navitoclax. (D) Mean (SD) senescence associated beta galactosidase activity in A549 epithelial cells when cultured with TSC2^−/−^ 621-101 cells, TSC2^+/+^ 621-103 cells and LAFs in combination and the effect of rapamycin, Dasatinb and Navitoclax. *p<0.05, **p<0.01, ****p<0.0001.

We examined the effect of the mTORC1 inhibitor rapamycin and the senolytic drugs Dasatinib (Src kinase inhibitor) and Navitoclax (Bcl-2, Bcl-XL and Bcl-w inhibitor) on SAβgal activity in combinations of *TSC2* null and addback cells, LAFs and lung epithelial derived A549 cells. In LAFs, the *TSC2* null dependent induction of SAβgal activity was inhibited by rapamycin but not Dasatinib or Navitoclax. In the epithelial cultures, rapamycin blocked SAβgal activity induced by the presence of *TSC2* null LAM cell co-cultures but the addition of LAFs to the A549/LAM 621-101 co-cultures induced SAβgal activity that was resistant to rapamycin. Dasatinib reduced epithelial SAβgal activity in the *TSC2* null 621-101/A549 co-cultures but had no effect upon the *TSC2* addback 621-103/LAF/A549 co-cultures (Figure 4C, D).

### LAM nodule cell interactions drive AT2 cell behaviour

To model these interactions using primary AT2 cells and examine the effect of LAM cell mTOR dysregulation on senescence induction *in vitro*, we made 3D LAM spheroid cultures (comprising co-cultures of either *TSC2* null 621-101 or *TSC2* add-back 621-103 cells with LAFs) which were then incubated with primary embryonic derived AT2 cell organoids (Figure 5A). Control AT2 organoids increased in size over 14 days, AT2 cell organoids co-cultured with individual LAF or *TSC2* addback 621-103 cells reduced organoid growth whereas *TSC2* null 621-101 cells induced rapid initial AT2 growth which plateaued early (Figure 5B). Both *TSC2* null and addback 621 cell / LAF spheroids tended to inhibit AT2 organoid growth over two weeks, although not significantly so (Figure 5C). To determine if this potential growth reduction was associated with the induction of senescence, organoids were immunostained for p16 and p21 proteins. Control organoids grown without LAM spheroids had low levels of expression of either protein. Both p16 and p21 were induced in the presence of *TSC2* null 621-101 containing spheroids, whereas *TSC2* addback 621-103 containing spheroids had a smaller and non-significant effect on p16 and p21 induction (Figure 5D&E).

**Figure 5.**
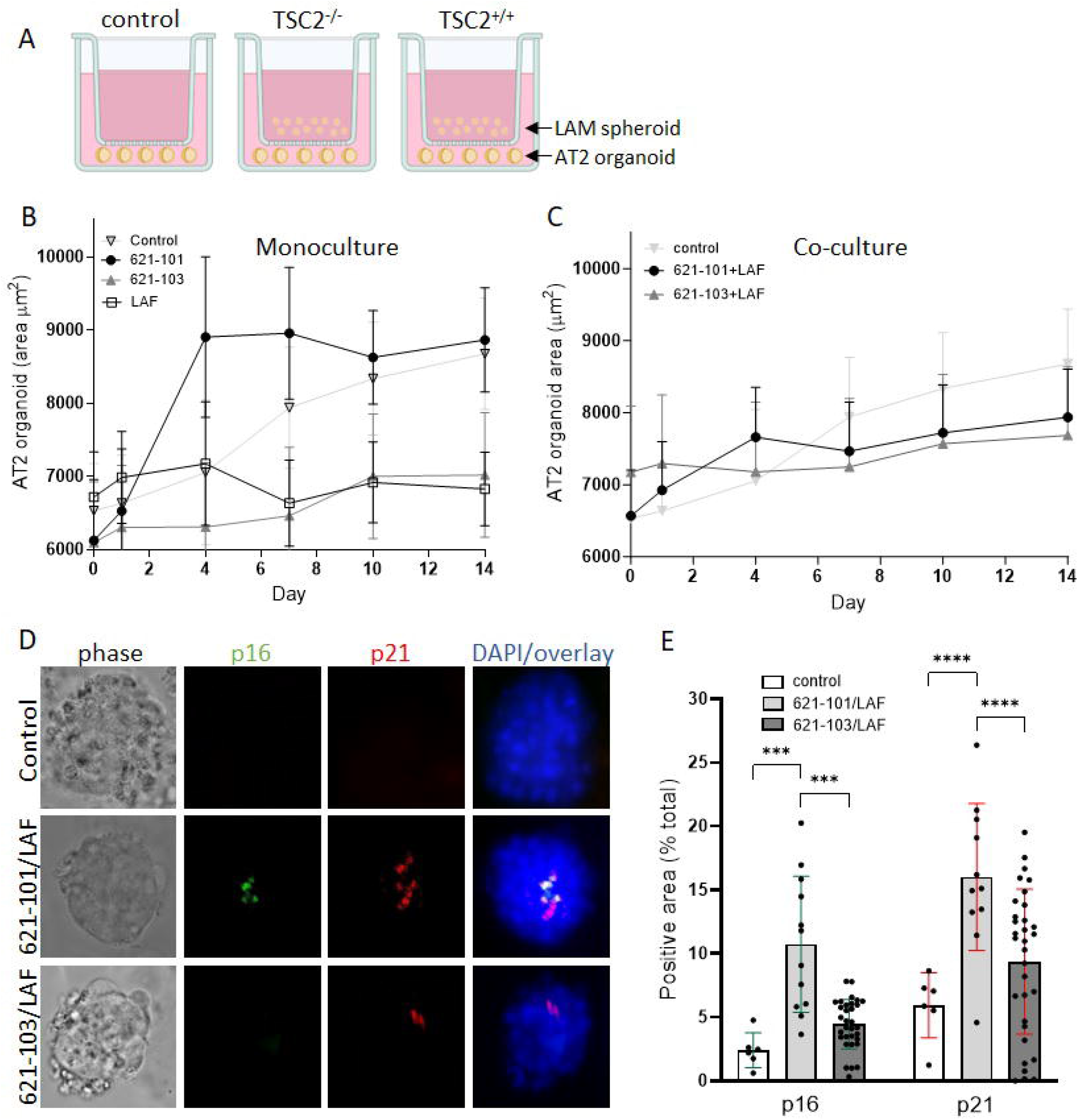
LAM nodules induce AT2 cell senescence *in vitro*. (A) Experimental setup showing LAM spheroids: 3D co-cultures of TSC2^−/−^ or TSC2^+/+^ LAM derived 621 cells and LAM associated fibroblasts (LAF) cultured in transwells with AT2 cell organoids over 14 days. (B) Mean (SD) AT2 cell organoid area over 14 days co cultured with TSC2^−/−^ 621-101 cells, TSC2^+/+^ 621-103 cells or LAFs. (C) Mean (SD) AT2 cell organoid area over 14 days co cultured with TSC2^−/−^ 621-101/LAF or TSC2^+/+^ 621-103/LAF co-cultures. (D) Induction of the senescence markers p16 and p21 in AT2 cell organoids over 14 days. (E) Quantification of p16 and p21 protein in AT2 cell organoids co-cultured with TSC2^−/−^ and TSC2^+/+^ LAM spheroids. ***p<0.001, ****p<0.0001.

### Inhibition of IL6 signalling facilitates epithelial wound repair processes and AT2 cell senescence

To understand the signalling events underlying AT2 cell senescence, scRNAseq data was used to examine the interactions between LAM nodules and AT2 cells. Upstream regulator analysis predicted activation of oestrogen receptor β (ESR2), the interleukin 6 receptor (IL6R), toll like receptor 7 (TLR7), TNF receptor superfamily 1A (TNFSF1A) and insulin dependent growth factor receptor 1 (IGF1R) (Figure 6A). Of importance to senescence induction, IL6R, TNFSF1A, IGF1R and ESR2 are upstream of *CDKN1A* / p21 (Table 2).

**Figure 6.**
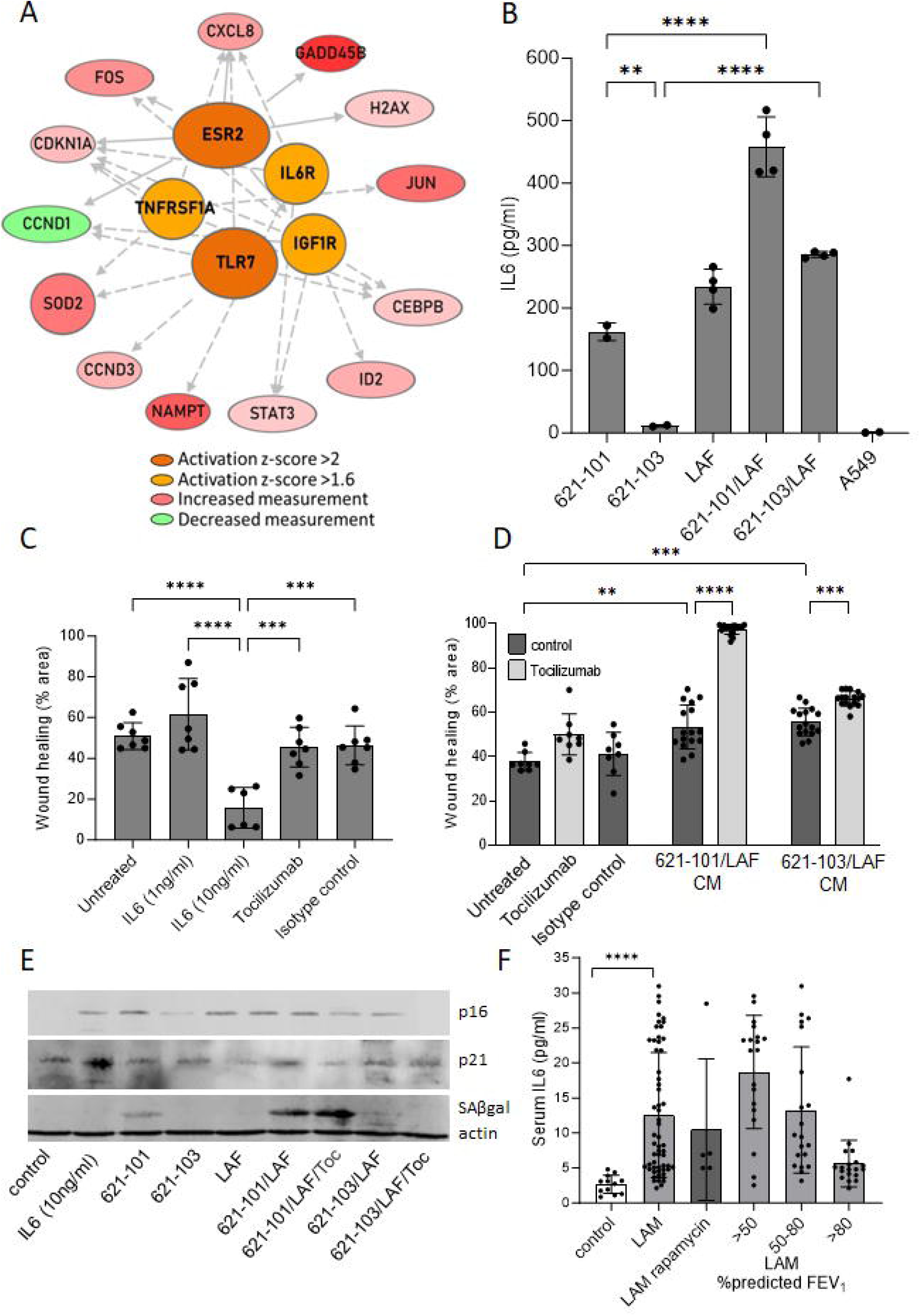
IL6 signalling induced by LAM nodules affects wound repair *in vitro*. (A) Upstream regulator analysis of AT2 cell gene induction predicted activation of oestrogen receptor β (ESR2), the interleukin 6 receptor (IL6R), toll like receptor 7 (TLR7), TNF receptor superfamily 1A (TNFSF1A) and insulin dependent growth factor receptor 1 (IGF1R). (B) Secretion of IL6 by TSC2^−/−^ 621-101 cells, TSC2^+/+^ 621-103 cells and LAM associated fibroblasts (LAF), LAM cell/LAF co-cultures and epithelial A549 cells. (C) The effect of IL6 on scratch wound repair in A549 cells. Graphs show mean (SD) percent of wound area healed over 24 hours. (D) The effect of IL6 inhibition using Tocilizumab on scratch wound repair in A549 epithelial cells incubated with conditioned medium (CM) from 3D co-cultures of TSC2^−/−^ 621-101 or TSC2^+/+^ 621-103 LAM derived 621 cells with LAM associated fibroblasts (LAF). (E) Western blots showing expression of p16, p21, and senescence associated β-galactosidase (SAβgal) protein expression by AT2 cell organoids over 14 days co-cultured with TSC2^−/−^ 621-101 cells, TSC2^+/+^ 621-103 cells and LAM associated fibroblasts (LAF) in mono or co-cultures as described and the effect of Tocilizumab (Toc). Beta actin is used as a loading control. (F) Serum IL6 in healthy control women, women with LAM untreated with an mTOR inhibitor and women with LAM treated with rapamycin. Stratification divides the LAM group according to their percent predicted forced expiratory volume in 1 second (FEV_1_) at the time of sampling. **p<0.01, ***p<0.001, ****p<0.0001.

**Table 2.**
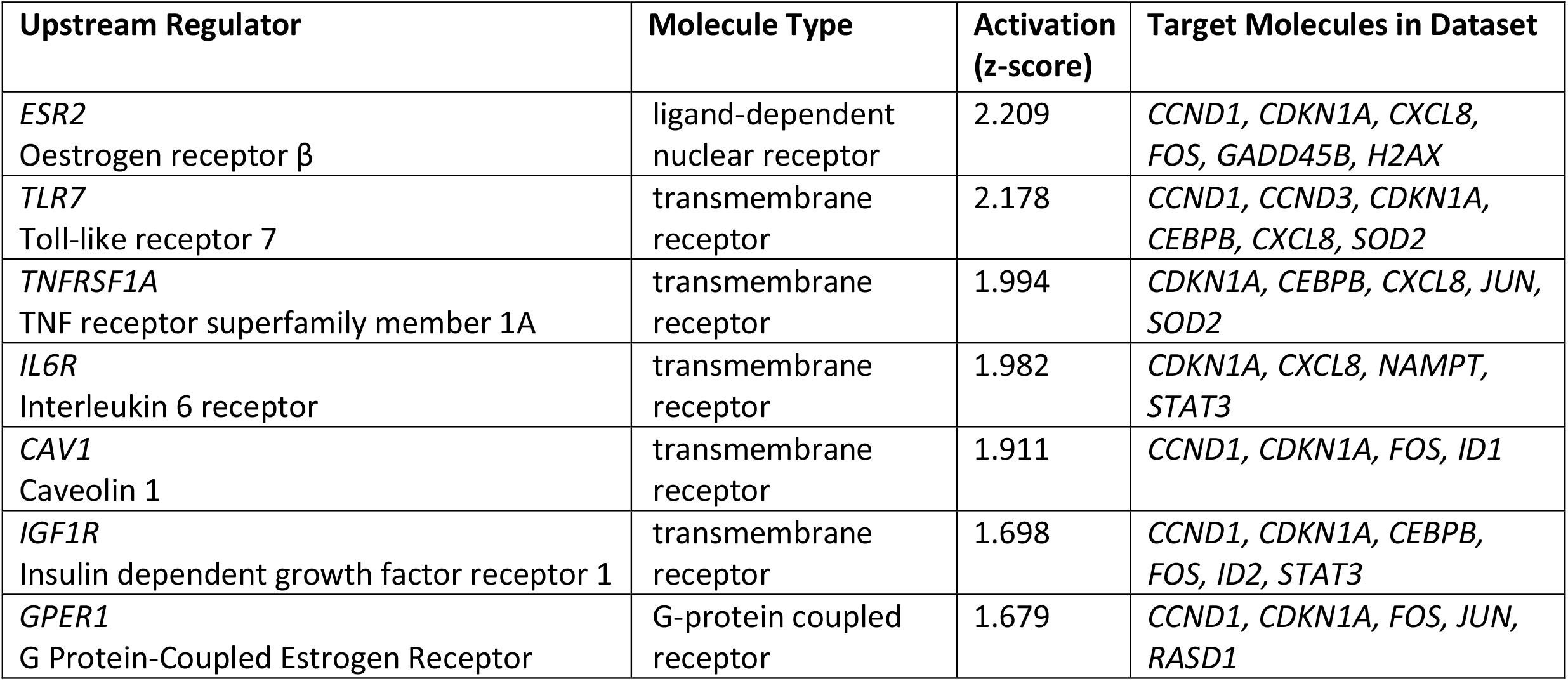
Predicted AT2 cell receptor activation from up-stream regulator analysis.

| Upstream Regulator | Molecule Type | Activation (z-score) | Target Molecules in Dataset |
| --- | --- | --- | --- |
| *ESR2*<br>Oestrogen receptor $\beta$ | ligand-dependent nuclear receptor | 2.209 | *CCND1, CDKN1A, CXCL8, FOS, GADD45B, H2AX* |
| *TLR7*<br>Toll-like receptor 7 | transmembrane receptor | 2.178 | *CCND1, CCND3, CDKN1A, CEBPB, CXCL8, SOD2* |
| *TNFRSF1A*<br>TNF receptor superfamily member 1A | transmembrane receptor | 1.994 | *CDKN1A, CEBPB, CXCL8, JUN, SOD2* |
| *IL6R*<br>Interleukin 6 receptor | transmembrane receptor | 1.982 | *CDKN1A, CXCL8, NAMPT, STAT3* |
| *CAV1*<br>Caveolin 1 | transmembrane receptor | 1.911 | *CCND1, CDKN1A, FOS, ID1* |
| *IGF1R*<br>Insulin dependent growth factor receptor 1 | transmembrane receptor | 1.698 | *CCND1, CDKN1A, CEBPB, FOS, ID2, STAT3* |
| *GPER1*<br>G Protein-Coupled Estrogen Receptor | G-protein coupled receptor | 1.679 | *CCND1, CDKN1A, FOS, JUN, RASD1* |

IL6 is a canonical SASP protein and activation of the IL6 receptor suggests its involvement in the senescence response in LAM. To investigate this further, we first examined IL6 protein secretion by LAM related cells. TSC2^−/−^ 621-101 cells produced IL6 whereas TSC2^+/+^ 621-103 cells expressed only minimal IL6. LAFs produced IL6 whereas lung epithelial derived A549 cells did not. Co-cultures of 621-101 cells and LAFs, matched for total cell number, produced more IL6 than either cell type alone (Figure 6B). To determine if IL6 signalling could impact on lung repair we treated scratch wounded alveolar epithelial A549 cells with recombinant IL6. IL6 (10ng/ml) significantly inhibited scratch wound resolution over 24 hours (Figure 6C). Repair of scratch wounded A549 cells incubated with serum-containing conditioned media from TSC2^−/−^ 621-101 cells / LAF spheroids was markedly enhanced by Tocilizumab which had a lesser but still significant effect on wound healing of TSC2^+/+^ 621-103 / LAF spheroid conditioned medium (Figure 6D).

To determine if IL6 could mediate senescence in AT2 cells, we examined if Tocilizumab would reduce the accumulation of p21 and p16 in AT2 cell organoids exposed to TSC2^−/−^ 621-101/LAF spheroids over 7 days. IL6 induced p16, and to a lesser extent p21 in AT2 cell organoids. 621-101 cells induced p16 and SAβgal whereas the 621-103 TSC2 addback cells had a smaller effect on p16 and did not induce SAβgal. 621-101/LAF co-cultures strongly induced SAβgal, p21 and p16. 621-103/LAF co-cultures had lesser effects on p21 and p16 induction and had no effect on SAβgal. Tocilizumab had only modest effects on p16 and p21 induction by 621-101/LAF co-cultures and did not affect SAβgal but supressed p16 induction by 621-103/LAF co-cultures (Figure 6E). Similar results were seen in A549 cells (Supplementary figure 5).

To determine if IL6 is increased in patients with LAM, we compared serum IL6 in healthy control women, 58 women with LAM untreated with an mTOR inhibitor and five women receiving rapamycin for LAM. There was a greater than four-fold elevation of serum IL6 in women with LAM (p=0.003). Stratifying patients according to their lung function, showed an inverse correlation between serum IL6 level and FEV_1_ tertiles (p<0.0001. Figure 6F). Although IL6 was lower in subjects treated with rapamycin than rapamycin naive women, the difference was not significant. Examining six-minute walk distance as a functional measure of patient performance, greater serum IL6 values were seen in those with the lowest six-minute walk distance (R^2^=0.14, p=0.046. Supplementary figure 6A). Surprisingly IL6 was not correlated with serum VEGF-D (R^2^ =0.0009, p=0.85), a well characterised LAM biomarker, thought to be related to mTOR dysregulation (Supplementary figure 6B).

## Discussion

We have shown that senescent cells accumulate in LAM nodules and lung parenchyma in a TSC2 / mTOR dependent manner. mTOR dysregulation drives secretion of the canonical SASP factor IL6 by LAM cells which contributes to the induction of senescence in AT2 cells and impairs epithelial wound healing. Further, the IL6 receptor antagonist Tocilizumab can enhance epithelial repair in the presence of LAM spheroids *in vitro*. Our findings point to a mechanism whereby senescence inhibits epithelial repair and increases the effect of lung injury in LAM and that inhibiting IL6 signalling in parallel with mTOR inhibition may reduce lung injury in LAM.

We observed that LAM lungs contained many more senescent cells than healthy lung tissue and hypothesised that cellular senescence would impair the repair process to proteolytic injury in LAM, thus accelerating tissue damage. Consistent with this idea, RNA sequencing showed increased activity in cytokine response pathways, consistent with SASP signalling and senescence in addition to injury as shown by increased apoptosis in lung parenchyma. scRNAseq data from LAM lungs showed that the SenMayo gene panel^26^, a set of 125 genes which identify senescent cells in multiple settings was increased in LAM when compared with control AT2 cells. Utilising scRNAseq from a patient treated with rapamycin, we observed that AT2 cell related senescent gene expression could be reduced by mTOR inhibitor therapy, reinforcing the role of mTOR dysregulation in LAM and AT2 cell senescence.

Senescence is a process of irreversible cell cycle arrest, thought to have evolved to suppress tumorigenesis in cells prone to DNA damage and can be induced by high levels of cell division, injurious processes including reactive oxygen species, DNA damage or oncogene activation including that resulting in mTOR dysregulation^22^. We had hypothesised that TSC2 loss/mTOR dysregulation would initially induce senescence in LAM cells. Consistent with this idea, scRNAseq data showed a doubling of senescence markers in LAM^CORE^ cells. *In vitro*, *TSC2* null cells express SAβgal, p16, p21 and IL6 at higher levels than their *TSC2* addback counterparts and in co-cultures LAM derived 621-101 cells and 621-101/LAF cultures increase expression of SAβgal in LAFs and alveolar epithelial cells to a greater extent than TSC2 addback 621-103 cells and 621-103/LAF co-cultures. In LAM lung tissue however, dual IHC suggested only a minority of the p16 or p21 expressing cells were LAM cells. We also observe that LAFs express senescence markers and can also increase SAβgal expression in LAM derived 621-101 cells which was not the case in the TSC2 addback 621-103 cells. Together these findings suggest that senescence is a feature of LAM lung tissue where mTOR dysregulation is necessary but further cell/cell interactions are likely to be required to generate the full senescent phenotype.

Senescence is a feature of other lung diseases, particularly those affecting older people where replicative senescence and consequent p53 activation are important in addition to reactive oxygen species generation in COPD and telomere dysfunction in lung fibrosis^19^ ^27^. In COPD, activation of mTOR signalling is associated with induction of MicroRNA-34a which suppresses sirtuin-1 to promote p53-induced senescence and an NF-κB, mediated SASP response^19^ ^28^. In COPD rapamycin both inhibits senescence and the SASP^29^ and whilst mTOR dysregulation can induce changes consistent with COPD in animals, unlike LAM, mTOR dysregulation in COPD is not likely to be the initiating event prompting senescence^29^. Telomere dysfunction, aging and other processes induce senescence in idiopathic pulmonary fibrosis (IPF), where senescent myofibroblasts induce senescence via SASP proteins in adjacent cells^30^. People with LAM are younger than those affected by COPD and IPF and also less likely to smoke^31^, our data suggest that in LAM, interactions between mTOR dysregulated LAM cells and LAFs suggest that cell-cell interactions within LAM nodules support the generation of senescent cells in an mTOR dependent manner, possibly less dependent upon aging, ROS production or telomere dysfunction seen in other lung diseases.

We used a number of methods to understand how cell-cell communication within the LAM lung leads to AT2 cell senescence which could adversely affect the response to lung injury. RNAseq of lung parenchymal cells highlighted activation of cytokine response genes, consistent with SASP signalling, defence against injury, apoptosis and senescence reflecting the LAM related parenchymal damage and the repair response to that injury. We focused upon the AT2 cell, the stem cell responsible for lung repair and prone to senescence^32^. Senescent AT2 cells were not detected in healthy lungs but present in the AT2 cells surrounding LAM nodules^25^, in cyst walls and in more normal appearing areas of LAM lung. scRNAseq of AT2 cells allowed us to examine the senescence response in more detail and observed a clear induction of the SenMayo gene set in LAM compared with healthy AT2 cells.

LAM spheroids induced p16 and p21 in human AT2 cell organoids, dependent on mTOR activation, consistent with SASP factors generated by senescent cells potentially affecting wound repair processes and thus contribute to lung cyst formation. Whilst it is difficult to relate senescence directly to wound repair in these models, the induction of these cell cycle inhibitors is likely to affect growth and we observed AT2 cell organoid growth was altered by the presence of both TSC2 null and TSC2 addback LAM spheroids. Whilst these *in vitro* changes over 14 days were not significant, it is possible that the markers of AT2 cell senescence we observed in human LAM tissue represents a similar process, which could enhance lung cyst formation over the disease course. Upstream regulator analysis highlighted the IL6 receptor as a potential effector of AT2 cell senescence. We showed that IL6 was produced by 621-101 cells in a TSC2 dependent manner which inhibited wound repair and induced AT2 cell SAβgal and p16 expression *in vitro*. Interestingly, although LAM spheroids did not alter scratch wound repair, treatment with the IL6R inhibitor Tocilizumab strongly enhanced wound repair suggesting IL6R signalling inhibits wound repair but other features of the LAM spheroid secretome have competing effects on these processes. The findings that IL6 is elevated in the serum of women with LAM^33^ and as we show here, related to the extent of lung involvement shown by lung function and patient performance, makes IL6 signalling an attractive potential target for therapy. Whilst some IL6 production is clearly a consequence of mTOR dysregulation and rapamycin sensitive, the production of IL6 by LAFs that do not have TSC loss and mTOR dysregulation and the lack of correlation between serum levels of IL6 and VEGF-D, thought to be a readout of pathological mTOR signalling in women with LAM, suggest that IL6 antagonism could be trialled in conjunction with mTOR inhibition.

Our findings using a range of techniques and senescence markers highlight that senescent cells accumulate in both LAM nodules and the AT2 cell population and moreover, these processes which include canonical SASP factors such as IL6 are mTOR dependent. In parallel to the induction of senescence, LAM nodule SASP factors alter epithelial cell growth and repair in co-cultures. Whilst it is difficult to definitively link senescence and lung repair, these processes are related and our findings highlight that targeting SASP factors including IL6 in parallel with mTOR inhibition, could potentially reduce lung damage in LAM.

## Methods

### Patient samples

Serum samples and lung tissue were obtained from patients receiving care at the UK LAM Centre between 2011 and 2022. All patients had LAM as defined by American Thoracic Society / Japanese Respiratory Society criteria^34^. Tissue samples taken for clinical care, either for diagnosis or from surgical treatment for pneumothorax were all from rapamycin naive patients. Linked clinical data was captured from medical notes and lung function (forced expired volume in on second, FEV_1_, lung diffusion of carbon monoxide, DL_CO_ and six-minute walk distance) and serum vascular endothelial growth factor were recorded at baseline and lung function repeated at subsequent visits. Change in lung function was calculated as the slope of a regression line of all values of FEV_1_ (ΔFEV_1_) or DL_CO_ (ΔDL_CO_). Clinical characteristics of patient samples used for laser capture microdissection and immunohistochemistry are shown in Table 1. The study was approved by the East Midlands Research Ethics Committee (reference 13/EM/0264) and all participants gave written informed consent.

### Cell, LAM spheroid and alveolar organoid culture

TSC2-null 621-101 and TSC2–add-back 621-103 cells were a gift from Dr. Lisa Henske (Brigham and Women’s Hospital). TTJ cells derived from renal tumours of TSC2^+/−^ C57BL/6 mice, were provided by Dr. Vera Krymskaya (University of Pennsylvania). Primary LAM associated fibroblasts were obtained from lung tissue from women with LAM and LAM spheroids comprising 621 cells and LAFs were prepared from 10,000 cells in ultra-low attachment 96-well plates as previously described^6^. Human foetal lung tissue was provided from terminations of pregnancy from the MRC/Wellcome Trust Human Developmental Biology Resource (London and Newcastle, University College London (UCL) site REC reference: 18/LO/0822; Project 200591; www.hdbr.org)^35^. Sample age ranged from 17-22 weeks of gestation determined by external physical appearance and measurements. Samples had no known genetic abnormalities. Sample gender was unknown at the time of collection and was not determined. AT2 cells were derived from this foetal material and were cultured in organoids as described^36^.

### Laser capture microdissection

Tissue sections were mounted on PEN Membrane Glass Slides (Thermo Fisher scientific LCM0522), deparaffinised in xylene for 2 minutes, twice, 100% ethanol for 1 minute, 96% ethanol for 1 minute then 70% ethanol for 1 minute and stained using Cresyl Violet. LCM was carried out using a Zeiss AxioImager Z1 (Zeiss, Thornwood, NY), using αSMA and PNL2 staining to identify LAM nodules. Microdissected samples were captured by AdhesiveCap 500ul microcentrifuge tubes (Fisher Scientific, Carl Zeiss™ 415190-9211-000). RNA extraction was performed by using RNeasy DSP FFPE Kit (QIAGEN, ID: 73604). Genomic DNA was eliminated using, RQ1 RNase-Free DNase (Promega, M6101) and clean-up performed using Monarch® RNA Cleanup Kit (Biolabs, New England, T2030L).

### Library Preparation and RNA Sequencing

Laser captured RNA samples were quantified using Ribogreen QuanT-iT kit on a Fluostar Plate reader and quality checked using a high sensitivity RNA Tape on a 4200 TapeStation. The RIN number was between 1.7 to 3.3. and the percentage of RNA molecules that are greater than 200bp in length (DV200 score) also calculated, with samples with higher than 35% DV200 chosen. The library preparation and RNA sequencing was performed by Birmingham Genomic centre using Lexogen QuantSeq 3’ mRNA-Seq Library Prep Kit FWD from Illumina. Libraries were quantified using a PicoGreen Quant-iT kit and sized with a D1000 tape. For sequencing, libraries were pooled in equal volumes according to their molarity. These were then quality checked using the Agilent TapeStation DNA 1000 tape and DNA HS Qubit. Pooled libraries was taken into sequencing at 2nM and loaded on the NovaSeq 6000 at 400pM with an SP 100 cycle flowcell.

### Single cell RNA sequencing analysis

LAM lung scRNAseq data were retrieved from the LAM Cell Atlas with scRNAseq data QC, pre-filtering, batch correction and data integration performed as previously described^37^. 65,287 cells from 11 LAM lungs (13 biological replicates) and 43,128 cells from 8 control female lungs^11^ ^38^ were included in integrative analysis using Seurat 3^39^. Unbiased cell clustering was performed using the Leiden algorithm^40^. Cell clusters were mapped to cell types based on expression of known cell type marker and signature genes from our previous LAM and normal lung single cell studies^37^ ^41^ ^42^. Automated cell type annotation was done using our previous LAM single cell study and our recently released LungMAP CellRef as references^37^ ^43^. Integrated single cell analysis identified an AT2 cluster consisting of 12,291 LAM AT2 cell and 17,378 control AT2 cells. The Wilcoxon rank-sum test was used to compare the gene expression levels between AT2 cells in LAM and control. The fold change (FC) was calculated as the ratio of the average expression level in the AT2 populations in LAM lungs and in normal control lungs. Functional enrichment analysis was performed using ToppGene suite^44^. CellChat^45^ analysis was applied on the scRNA-seq data to decipher LAM^CORE^ and LAM AT2 cell communication patterns based on cell type selective expression of ligands secreted from LAM^CORE^ cells and receptors expressed on the surface of LAM AT2 cells. Ingenuity Pathway Analysis (IPA) was used to predict activated upstream regulators using the differentially expressed senescence genes in LAM AT2 cells as input genes.

### Bulk RNA-seq analysis

RNA sequencing analyses (RNA-seq) were performed on laser capture microdissection of LAM nodule and the rest of the lung as self-control. RNA-seq data alignment, QC and normalization was done using the RNA-Seq workflow in Partek Flow software (v7. Partek Inc., St. Louis, MO, USA). Human genome hg38 was used as reference. Gene counts were normalized using the median of ratios (i.e., ratio of read count to its geometric mean across all samples before being subjected to a differential expression analysis^46^. Differentially expressed genes (DEGs) were identified using DESeq2 in Partek with the combination criteria of a p-value < 0.05 and absolute fold change >1.5. Genes with mean average count across all samples < 10 were removed for further analysis. DESeq2 was used to find differentially expressed genes between the laser capture samples and their corresponding rest of lung samples for each clinical measurement group.

### Immunohistochemistry

Tissue for immunohistochemistry was obtained from 20 biopsies and four explanted lungs from transplantation and three control female lungs matched for age. Paraffin-embedded lung tissues were dewaxed in Xylene (Merck 214736, UK) and rehydrated with ethanol (100%, 70% each twice, and water). Antigen retrieval was carried out by heating sections sodium citrate buffer or Tris-EDTA buffer follow by quenching endogenous peroxidase activity by 3% H2O2. Primary antibodies used were: anti-melanoma associated antigen PNL2 (1:100, Zytomed MSK082-05), mouse monoclonal anti-smooth muscle actin (1:5000 Sigma Clone 1A4, A2547), mouse anti-Ki67 8D5 (1:250, Cell Signaling Technology, No. 9449), P16^INK4A^ Rabbit Polyclonal antibody (1:500, Proteintech No. 10883-1-AP), p21^Waf1/Cip1^ (12D1) Rabbit mAb (1:50 Cell Signaling Technology No: 2947), anti-pro-surfactant protein C antibody Rabbit monoclonal (1:1,000, Abcam, ab90716). Secondary antibodies used were horseradish peroxidase (HRP)-conjugated goat anti-rabbit or anti-mouse (ImmPRESS, Vector Laboratories, MP-7452). Detection of primary antibodies was performed using ImmPACT 3,3′-diaminobenzidine (DAB) peroxidase substrate (Vector Laboratories, SK-4105). Sections were counterstained with Mayer’s hematoxylin. Fluorescence secondary antibodies used were Goat anti-Rabbit IgG, Alexa Fluor™ 488 (Invitrogen, A-11008), Goat anti-Mouse IgG, Alexa Fluor™ 488 (Invitrogen A-11001), Donkey anti-Rabbit Antibody, Alexa Fluor™ 594 (Invitrogen, A-21207) and Donkey anti-Mouse IgG, Alexa Fluor™ 594 (Invitrogen, A-21203).

### Immunofluorescence

Organoids were collected after centrifugation at 500G for 5 minutes. Samples were fixed with either 4% paraformaldehyde (PFA) or ethanol based on the primary antibody protocol for 30 min at RT, follow by permeabilization using 0.1% Triton X-100, 0.05% Tween-20 in PBS. Samples were blocked in 3% BSA in PBS. Primary antibodies were incubated at 4 °C overnight and secondary antibodies incubated for 1 hour room temperature and counterstained with DAPI. AT2 Organoids were fixed with PFA for 3 minutes and then released from matrigel by incubating the culture with Corning Cell Recovery Solution (Merck, CLS354253) for 2 hours at 4 °C.

### Senescence associated β-galactosidase activity

Senescence associated β-galactosidase was measured in cell cultures using a plate-based fluorescent assay (Cell Signaling, 23833). In tissues β-galactosidase activity at pH 6 was determined using either Senescence β-Galactosidase Staining Kit (cell signaling, 9860) or the CellEvent Senescence Green Detection Kit (Invitrogen, C10850) according to the conditions.

### Animal modelling

TTJ cells (10^6^ cells in 150ml of PBS) were injected into the tail vein of B6 Albino mice (B6N-Tyrc-Brd/BrdCrCrl) with sham injections (150ml of PBS) used for control animals^6^. After the TTJ cell–injected untreated group had lost 10% of body weight, the study was terminated in all groups and senescence associated β-galactosidase measured in lung tissue. The project was conducted under Home Office project Personal Project Licence P435A9CF8.

### Western blotting

Nuclear and cytoplasmic protein fractions were separated using a subcellular fractionation protocol as described^47^. Blots were probed with primary antibodies overnight and secondary antibodies for one hour room temperature. Primary antibody P16-INK4A Rabbit Polyclonal antibody (1:100, Proteintech No. 10883-1-AP), p21 Waf1/Cip1 (12D1) Rabbit mAb (1:100 Cell Signaling Technology No: 2947) and β-actin Rabbit mAb (Cell signaling, 4970) was used as a loading control. Clarity Max Western ECL Substrate was used for HRP conjugated (BIO-RAD 1705062) for detecting the protein bands.

### IL6 ELISA

Conditioned media were passed through a 0.2μM filter and IL-6 quantified using a Human IL-6 DuoSet ELISA (R and D, DY206) according to the manufacturer’s protocol. The optical density measured at 450 nm with wavelength correction of 540 nm. Serum IL6 was determined using the same kit in 66 LAM patients and 13 control subjects.

### Data analysis

Data were analysed by Mann-Whitney test, and two-way ANOVA using the Dunnett multiple comparisons method as appropriate using Graph Pad Prism 10 (GraphPad Software Boston, MA). The correlation of P16 and p12 positive cells with future lung function change, adjusted for baseline lung function, was analysed in PRISM. P < 0.05 was considered as indicating significance.

## Data availability

LAM lung RNA sequencing data are available at GEO (GSE265851). LAM lung single cell data is available through the LAM cell atlas (https://research.cchmc.org/pbge/lunggens/LCA/LCA.html). Link anonymised clinical data are available to academic groups through the corresponding author.

## Author contributions

RB-J, DC and SM, performed the experimental work. YW, KC and YX performed the bioinformatic analysis. MP and RC helped with laser capture microdissection, KL and ELR contributed AT2 cells and advised on the work. SRJ conceived and directed the study, obtained the funding and saw the patients. RB-J and SRJ wrote the manuscript with all authors providing intellectual input and approving the final version.

## References

1. McCarthy C, Gupta N, Johnson SR, et al. Lymphangioleiomyomatosis: pathogenesis, clinical features, diagnosis, and management. Lancet Respir Med 2021 doi: 10.1016/s2213-2600(21)00228-9 [published Online First: 2021/08/31]

2. Seibert D, Hong C-H, Takeuchi F, et al. Recognition of Tuberous Sclerosis in Adult Women: Delayed Presentation With Life-Threatening Consequences. Annals of Internal Medicine 2011;154(12):806–13. doi: 10.1059/0003-4819-154-12-201106210-00008

3. Clements D, Dongre A, Krymskyaya V, et al. Wild Type Mesenchymal Cells Contribute to the Lung Pathology of Lymphangioleiomyomatosis. PLoS ONE 2015 doi: DOI: 10.1371/journal.pone.0126025

4. Kumasaka T, Seyama K, Mitani K, et al. Lymphangiogenesis in Lymphangioleiomyomatosis: Its Implication in the Progression of Lymphangioleiomyomatosis. Am J Surg Pathol 2004;28(8):1007–16.

5. Liu HJ, Lizotte PH, Du H, et al. TSC2-deficient tumors have evidence of T cell exhaustion and respond to anti-PD-1/anti-CTLA-4 immunotherapy. JCI insight 2018;3(8) doi: 10.1172/jci.insight.98674 [published Online First: 2018/04/20]

6. Babaei-Jadidi R, Dongre A, Miller S, et al. Mast Cell Tryptase Release Contributes to Disease Progression in Lymphangioleiomyomatosis. Am J Respir Crit Care Med 2021 doi: 10.1164/rccm.202007-2854OC [published Online First: 2021/04/22]

7. Miller S, Stewart ID, Clements D, et al. Evolution of lung pathology in lymphangioleiomyomatosis: associations with disease course and treatment response. The journal of pathology Clinical research 2020 doi: 10.1002/cjp2.162 [published Online First: 2020/05/01]

8. Dongre A, Clements D, Fisher AJ, et al. Cathepsin K in Lymphangioleiomyomatosis: LAM Cell–Fibroblast Interactions Enhance Protease Activity by Extracellular Acidification. The American Journal of Pathology 2017;187(8):1750–62. doi: 10.1016/j.ajpath.2017.04.014

9. Hayashi T, Fleming MV, Stetler-Stevenson WG, et al. Immunohistochemical study of matrix metalloproteinases (MMPs) and their tissue inhibitors (TIMPs) in pulmonary lymphangioleiomyomatosis (LAM). Human pathology 1997;28(9):1071–8.

10. Zhe X, Yang Y, Schuger L. Imbalanced Plasminogen System in Lymphangioleiomyomatosis: Potential Role of Serum Response Factor. Am J Respir Cell Mol Biol 2004:2004–0289OC.

11. Obraztsova K, Basil MC, Rue R, et al. mTORC1 activation in lung mesenchyme drives sex- and age-dependent pulmonary structure and function decline. Nat Commun 2020;11(1):5640. doi: 10.1038/s41467-020-18979-4 [published Online First: 2020/11/08]

12. Hogan BL, Barkauskas CE, Chapman HA, et al. Repair and regeneration of the respiratory system: complexity, plasticity, and mechanisms of lung stem cell function. Cell Stem Cell 2014;15(2):123–38. doi: 10.1016/j.stem.2014.07.012 [published Online First: 2014/08/12]

13. Tan SY, Krasnow MA. Developmental origin of lung macrophage diversity. Development 2016;143(8):1318–27. doi: 10.1242/dev.129122 [published Online First: 2016/03/10]

14. Nabhan AN, Brownfield DG, Harbury PB, et al. Single-cell Wnt signaling niches maintain stemness of alveolar type 2 cells. Science 2018;359(6380):1118-23. doi: 10.1126/science.aam6603 [published Online First: 2018/02/09]

15. Zepp JA, Zacharias WJ, Frank DB, et al. Distinct Mesenchymal Lineages and Niches Promote Epithelial Self-Renewal and Myofibrogenesis in the Lung. Cell 2017;170(6):1134–48.e10. doi: 10.1016/j.cell.2017.07.034

16. Clements D, Miller S, Jadidi R, et al. Cross-talk between LAM cells and fibroblasts may influence alveolar epithelial cell behavior in Lymphangioleiomyomatosis. American Journal of Physiology-Lung Cellular and Molecular Physiology 2021;322 doi: 10.1152/ajplung.00351.2021

17. Shiraishi K, Nakajima T, Shichino S, et al. In vitro expansion of endogenous human alveolar epithelial type II cells in fibroblast-free spheroid culture. Biochem Biophys Res Commun 2019;515(4):579–85. doi: 10.1016/j.bbrc.2019.05.187 [published Online First: 2019/06/11]

18. Mercado N, Ito K, Barnes PJ. Accelerated ageing of the lung in COPD: new concepts. Thorax 2015;70(5):482.

19. Barnes PJ, Baker J, Donnelly LE. Cellular senescence as a mechanism and target in chronic lung diseases. American Journal of Respiratory and Critical Care Medicine 2019;200 doi: 10.1164/rccm.201810-1975TR

20. Antikainen H, Driscoll M, Haspel G, et al. TOR-mediated regulation of metabolism in aging. Aging Cell 2017;16(6):1219–33. doi: 10.1111/acel.12689

21. Hoare M, Narita M. Transmitting senescence to the cell neighbourhood. Nat Cell Biol 2013;15(8):887–9. doi: 10.1038/ncb2811 [published Online First: 2013/08/03]

22. Chaib S, Tchkonia T, Kirkland JL. Cellular senescence and senolytics: the path to the clinic. Nature Medicine 2022;28(8):1556–68. doi: 10.1038/s41591-022-01923-y

23. Laberge R-M, Sun Y, Orjalo AV, et al. MTOR regulates the pro-tumorigenic senescence-associated secretory phenotype by promoting IL1A translation. Nat Cell Biol 2015;17(8):1049–61. doi: 10.1038/ncb3195 http://www.nature.com/ncb/journal/v17/n8/abs/ncb3195.html#supplementary-information

24. Du Y, Guo M, Wu Y, et al. Lymphangioleiomyomatosis (LAM) Cell Atlas. Thorax 2023;78(1):85–87. doi: 10.1136/thoraxjnl-2022-218772 [published Online First: 2023/01/05]

25. Matsui K, W KR, Hilbert SL, et al. Hyperplasia of type II pneumocytes in pulmonary lymphangioleiomyomatosis. Arch Pathol Lab Med 2000;124(11):1642–8.

26. Saul D, Kosinsky RL, Atkinson EJ, et al. A New Gene Set Identifies Senescent Cells and Predicts Senescence-Associated Pathways Across Tissues. bioRxiv 2021:2021.12.10.472095. doi: 10.1101/2021.12.10.472095

27. Wrench CL, Baker JR, Monkley S, et al. Small airway fibroblasts from Chronic Obstructive Pulmonary Disease patients exhibit cellular senescence. American Journal of Physiology-Lung Cellular and Molecular Physiology;0(0):null. doi: 10.1152/ajplung.00419.2022

28. Baker JR, Vuppusetty C, Colley T, et al. MicroRNA-570 is a novel regulator of cellular senescence and inflammaging. Faseb j 2019;33(2):1605–16. doi: 10.1096/fj.201800965R [published Online First: 2018/08/30]

29. Houssaini A, Breau M, Kebe K, et al. mTOR pathway activation drives lung cell senescence and emphysema. JCI insight 2018;3(3) doi: 10.1172/jci.insight.93203

30. Schafer MJ, White TA, Iijima K, et al. Cellular senescence mediates fibrotic pulmonary disease. Nat Commun 2017;8:14532. doi: 10.1038/ncomms14532 [published Online First: 2017/02/24]

31. Ryu JH, Moss J, Beck GJ, et al. The NHLBI Lymphangioleiomyomatosis Registry: Characteristics of 230 Patients at Enrollment. Am J Respir Crit Care Med 2006;173(1):105–11.

32. Parimon T, Chen P, Stripp BR, et al. Senescence of alveolar epithelial progenitor cells: a critical driver of lung fibrosis. American Journal of Physiology-Cell Physiology 2023;325(2):C483–C95. doi: 10.1152/ajpcell.00239.2023

33. Wang J, Filippakis H, Hougard T, et al. Interleukin-6 mediates PSAT1 expression and serine metabolism in TSC2-deficient cells. Proc Natl Acad Sci U S A 2021;118(39) doi: 10.1073/pnas.2101268118 [published Online First: 2021/09/22]

34. McCormack FX, Gupta N, Finlay GR, et al. Official American Thoracic Society/Japanese Respiratory Society Clinical Practice Guidelines: Lymphangioleiomyomatosis Diagnosis and Management. American Journal of Respiratory and Critical Care Medicine 2016;194(6):748–61. doi: 10.1164/rccm.201607-1384ST

35. Kyungtae L, Eimear NR, Dawei S, et al. A novel human fetal lung-derived alveolar organoid model reveals mechanisms of surfactant protein C maturation relevant to interstitial lung disease. bioRxiv 2023:2023.08.30.555522. doi: 10.1101/2023.08.30.555522

36. Lim K, Donovan APA, Tang W, et al. Organoid modeling of human fetal lung alveolar development reveals mechanisms of cell fate patterning and neonatal respiratory disease. Cell Stem Cell 2023;30(1):20–37.e9. doi: 10.1016/j.stem.2022.11.013

37. Guo M, Yu JJ, Perl AK, et al. Single-Cell Transcriptomic Analysis Identifies a Unique Pulmonary Lymphangioleiomyomatosis Cell. American journal of respiratory and critical care medicine 2020;202(10):1373–87. doi: 10.1164/rccm.201912-2445OC

38. Reyfman PA, Walter JM, Joshi N, et al. Single-Cell Transcriptomic Analysis of Human Lung Provides Insights into the Pathobiology of Pulmonary Fibrosis. American journal of respiratory and critical care medicine 2019;199(12):1517–36. doi: 10.1164/rccm.201712-2410OC [published Online First: 2018/12/18]

39. Stuart T, Butler A, Hoffman P, et al. Comprehensive Integration of Single-Cell Data. Cell 2019;177(7):1888–902 e21. doi: 10.1016/j.cell.2019.05.031 [published Online First: 2019/06/11]

40. Traag VA, Waltman L, van Eck NJ. From Louvain to Leiden: guaranteeing well-connected communities. Sci Rep 2019;9(1):5233. doi: 10.1038/s41598-019-41695-z [published Online First: 2019/03/28]

41. Wang A, Chiou J, Poirion OB, et al. Single-cell multiomic profiling of human lungs reveals cell-type-specific and age-dynamic control of SARS-CoV2 host genes. Elife 2020;9 doi: 10.7554/eLife.62522 [published Online First: 2020/11/10]

42. Guo M, Du Y, Gokey JJ, et al. Single cell RNA analysis identifies cellular heterogeneity and adaptive responses of the lung at birth. Nat Commun 2019;10(1):37. doi: 10.1038/s41467-018-07770-1 [published Online First: 2019/01/04]

43. Guo M, Morley MP, Jiang C, et al. Guided construction of single cell reference for human and mouse lung. Nat Commun 2023;14(1):4566. doi: 10.1038/s41467-023-40173-5 [published Online First: 20230729]

44. Chen J, Bardes EE, Aronow BJ, et al. ToppGene Suite for gene list enrichment analysis and candidate gene prioritization. Nucleic Acids Res 2009;37(Web Server issue):W305–11. doi: 10.1093/nar/gkp427 [published Online First: 20090522]

45. Jin S, Guerrero-Juarez CF, Zhang L, et al. Inference and analysis of cell-cell communication using CellChat. Nat Commun 2021;12(1):1088. doi: 10.1038/s41467-021-21246-9 [published Online First: 2021/02/19]

46. Love MI, Huber W, Anders S. Moderated estimation of fold change and dispersion for RNA-seq data with DESeq2. Genome Biol 2014;15(12):550. doi: 10.1186/s13059-014-0550-8

47. Clements D, Mayer RJ, Johnson SR. Subcellular distribution of the TSC2 gene product tuberin in human airway smooth muscle cells is driven by multiple localization sequences and is cell-cycle dependent. Am J Physiol Lung Cell Mol Physiol 2007;292(1):L258–66.

48. Bankhead P, Loughrey MB, Fernández JA, et al. QuPath: Open source software for digital pathology image analysis. Scientific reports 2017;7(1):16878. doi: 10.1038/s41598-017-17204-5

