## Supplemental for "mTOR dysregulation induces IL6 and paracrine AT2 cell senescence impeding lung repair in lymphangioleiomyomatosis"

Supplementary tables and figures

**Supplementary table 1.** 100 most upregulated genes assessed by bulk RNA sequencing in laser capture isolated LAM nodules from 19 patients with LAM compared with normal lung tissue. The full dataset is available xxxxxxxxxxxxxxxx

| **Gene symbol** | **P-value (LAM vs. Control)** | **Fold change (LAM vs. Control)** | **LSMean (LAM) (LAM vs. Control)** | **LSMean (Control) (LAM vs. Control)** |
|---|---|---|---|---|
| *MMP11* | 0.009964729 | 269.2890793 | 269.2890793 | 1 |
| *ZNF324* | 0.002618641 | 156.1631409 | 156.1631409 | 1 |
| *ESRP2* | 0.013179556 | 125.7658885 | 182.5727108 | 1.451687043 |
| *MRGPRF* | 0.002686885 | 115.4989799 | 115.4989799 | 1 |
| *B3GNT8* | 0.046429956 | 97.10326966 | 97.10326966 | 1 |
| *TRIM63* | 0.026479265 | 94.01686195 | 151.8733889 | 1.615384579 |
| *FN3KRP* | 0.009515228 | 88.32529251 | 88.32529251 | 1 |
| *SLC18B1* | 0.01446082 | 85.56842825 | 85.56842825 | 1 |
| *ZBTB47* | 4.58475E-06 | 85.18364266 | 203.1302279 | 2.384615421 |
| *ARIH2OS* | 0.04832264 | 85.09948858 | 85.09948858 | 1 |
| *PDE6B* | 0.021687956 | 82.67230709 | 82.67230709 | 1 |
| *KLHL6* | 0.03239884 | 79.88639557 | 79.88639557 | 1 |
| *TTC39A* | 0.034483036 | 79.08945967 | 79.08945967 | 1 |
| *KIF21B* | 0.014195162 | 77.25483045 | 77.25483045 | 1 |
| *ABCB5* | 0.047272775 | 77.06538613 | 77.06538613 | 1 |
| *MINPP1* | 0.018562675 | 76.83336032 | 76.83336032 | 1 |
| *TRPV4* | 0.029434481 | 75.75740846 | 75.75740846 | 1 |
| *TMEM273* | 0.046352612 | 74.445846 | 74.445846 | 1 |
| *AGRP* | 0.009042088 | 74.1212295 | 74.1212295 | 1 |
| *WASH8P* | 0.005883626 | 73.82564502 | 111.153656 | 1.505623905 |
| *TFEB* | 0.044180349 | 70.01291569 | 70.01291569 | 1 |
| *HBP1* | 0.044560805 | 69.68997832 | 104.5385331 | 1.500051164 |
| *LY96* | 0.029953598 | 67.21379853 | 67.21379853 | 1 |
| *ZNF124* | 0.004484127 | 66.91309454 | 66.91309454 | 1 |
| *ITGBL1* | 0.016009334 | 66.07936179 | 66.07936179 | 1 |
| *CACNA1C* | 0.006254952 | 65.04068262 | 265.1658599 | 4.076923077 |
| *COL15A1* | 0.037650534 | 64.33278108 | 103.9221825 | 1.615384579 |
| *ABCC3* | 0.000286535 | 61.73773608 | 147.2207575 | 2.384615421 |
| *TMX3* | 0.008167493 | 58.75177363 | 92.35557856 | 1.571962391 |
| *TMEM254* | 0.012666949 | 54.67523868 | 88.32153741 | 1.615384579 |
| *HERPUD2* | 0.004503543 | 54.11452936 | 364.2324251 | 6.730769524 |
| *ZNF468* | 0.021826726 | 53.41839874 | 86.29125754 | 1.615384579 |
| *PUSL1* | 0.010066043 | 52.65842065 | 111.9494087 | 2.125954545 |
| *WDR46* | 0.029532138 | 52.09170221 | 52.09170221 | 1 |
| *BTBD3* | 0.043465562 | 49.86581414 | 49.86581414 | 1 |
| *MMP7* | 0.02045438 | 48.35176944 | 48.35176944 | 1 |
| *PSD4* | 0.002114871 | 48.28541983 | 232.3417809 | 4.811841373 |
| *PTPN22* | 0.045416904 | 48.1705179 | 77.81391177 | 1.615384579 |

| | | | | |
|---|---|---|---|---|
| *USP31* | 0.032651053 | 47.93020929 | 77.42572095 | 1.615384579 |
| *FAM13B* | 0.028659848 | 47.43003036 | 47.43003036 | 1 |
| *SAMD4A* | 0.043305301 | 45.7661526 | 45.7661526 | 1 |
| *EEPD1* | 0.045044065 | 44.43364912 | 44.43364912 | 1 |
| *NIPSNAP3A* | 0.002418215 | 43.38702904 | 68.2027779 | 1.571962391 |
| *REEP1* | 0.036820016 | 42.91731216 | 42.91731216 | 1 |
| *MYBBP1A* | 0.037644631 | 42.18050104 | 61.64842344 | 1.461538434 |
| *RAB4A* | 0.014934588 | 41.66483349 | 67.3047295 | 1.615384579 |
| *TUBA3FP* | 0.043446582 | 41.40106778 | 41.40106778 | 1 |
| *POP4* | 0.033767083 | 41.28777134 | 41.28777134 | 1 |
| *SLAMF1* | 0.02971143 | 41.23838938 | 66.61585825 | 1.615384579 |
| *C2orf74* | 0.029066569 | 41.09145329 | 132.7570028 | 3.230769228 |
| *DNAJC13* | 0.042834999 | 40.01053265 | 40.01053265 | 1 |
| *NUDCD2* | 0.004887862 | 39.87728419 | 83.69774275 | 2.098882721 |
| *ZNF526* | 0.023840127 | 38.75067583 | 60.91460504 | 1.571962391 |
| *NDFIP2* | 0.024414455 | 38.35969617 | 82.62088266 | 2.153846117 |
| *FNIP1* | 0.00152943 | 37.38129118 | 80.36950795 | 2.149992828 |
| *MIR4458HG* | 0.02121217 | 36.43005711 | 36.43005711 | 1 |
| *PAIP1* | 0.009413994 | 36.0556155 | 97.07284177 | 2.692308547 |
| *MTFP1* | 0.044782069 | 35.6642344 | 57.61145426 | 1.615384579 |
| *CASK* | 0.018776248 | 35.35325218 | 35.35325218 | 1 |
| *UBQLN2* | 0.047269592 | 34.63151664 | 34.63151664 | 1 |
| *RRN3* | 0.001842464 | 34.27916462 | 73.83204562 | 2.153846117 |
| *ZNF780B* | 0.038688513 | 33.96796958 | 33.96796958 | 1 |
| *RASGRF1* | 0.024198664 | 33.23967821 | 71.59315185 | 2.153846117 |
| *ANKS1B* | 0.030157088 | 32.87523363 | 32.87523363 | 1 |
| *AZIN1* | 2.86268E-05 | 32.69527222 | 96.82830605 | 2.961538458 |
| *PRPS2* | 0.014718588 | 32.18198898 | 51.98628872 | 1.615384579 |
| *MPI* | 0.00526314 | 31.84420849 | 144.2727916 | 4.530581804 |
| *CPED1* | 0.004182107 | 31.61256011 | 51.0664421 | 1.615384579 |
| *BAG3* | 0.018451605 | 31.3730299 | 31.3730299 | 1 |
| *MOCOS* | 0.035783658 | 30.14709246 | 48.69914826 | 1.615384579 |
| *GMCL1* | 0.000880407 | 30.01472068 | 202.0221673 | 6.730769524 |
| *COA4* | 0.001196928 | 29.97050652 | 80.68985087 | 2.692308547 |
| *PRKCA* | 0.009781408 | 28.9454801 | 77.93016348 | 2.692308547 |
| *HERC2* | 0.000483275 | 28.77297612 | 85.21227532 | 2.961538458 |
| *LRP2* | 0.047726953 | 27.60730878 | 27.60730878 | 1 |
| *CBWD3* | 0.034784119 | 27.56930914 | 42.23065401 | 1.531799502 |
| *CREG1* | 0.0013251 | 27.46807189 | 132.1720048 | 4.811841373 |
| *ZNF330* | 0.016750249 | 27.20878204 | 58.60352955 | 2.153846117 |
| *SCAI* | 0.014609859 | 27.04236583 | 190.3371349 | 7.038479402 |
| *COX18* | 0.043693342 | 26.42418025 | 78.25622603 | 2.961538458 |
| *BRCC3* | 0.03302363 | 25.79044039 | 25.79044039 | 1 |
| *CENPC* | 0.049512851 | 25.51200085 | 60.83631067 | 2.384615421 |

| | | | | |
|---|---|---|---|---|
| *KCNN4* | 0.020488461 | 25.43452669 | 54.78205654 | 2.153846117 |
| *LOC101930085* | 0.04864007 | 24.65906918 | 39.83388009 | 1.615384579 |
| *MRPS22* | 0.020083362 | 24.58514622 | 24.58514622 | 1 |
| *MEPCE* | 0.01570748 | 24.05256965 | 215.1090363 | 8.943287116 |
| *TMEM131* | 0.043581672 | 23.91394658 | 51.50696098 | 2.153846117 |
| *MFSD1* | 0.020894764 | 23.61842752 | 76.30568886 | 3.230769228 |
| *DERL1* | 0.001134589 | 23.39209571 | 101.6656416 | 4.346153626 |
| *NOM1* | 0.033069138 | 22.78797241 | 54.34055043 | 2.384615421 |
| *SRRD* | 0.043624123 | 22.48154231 | 33.72976381 | 1.500331398 |
| *TXNRD2* | 0.041359093 | 22.43136662 | 74.22392505 | 3.308934597 |
| *AP1S3* | 0.000821575 | 22.26630472 | 705.0110887 | 31.66268932 |
| *NSUN3* | 0.023614577 | 21.79710146 | 83.83500482 | 3.846153809 |
| *NTM* | 0.013592975 | 21.34528967 | 701.0215481 | 32.84197868 |
| *SH3PXD2B* | 0.027826486 | 21.22578253 | 142.8658501 | 6.730769524 |
| *VIRMA* | 0.005658337 | 20.89635237 | 61.88535117 | 2.961538458 |
| *URB1* | 0.030636798 | 20.8855265 | 49.14393648 | 2.353014011 |
| *NUFIP1* | 0.04922981 | 20.10746966 | 32.4812964 | 1.615384579 |
| *BRD8* | 0.003233504 | 19.70280121 | 68.11473792 | 3.457109332 |

**Supplementary table 2.** AT2 cell senescence-related genes from the SenMayo panel up-regulated in LAM.

| **Gene** | **P value** | **Fold change** (average log2) | **LAM** | **Control** |
|---|---|---|---|---|
| CCL20 | 8.26E-221 | 1.3586 | 0.227 | 0.099 |
| NAMPT | 0 | 1.268706 | 0.718 | 0.539 |
| CXCL2 | 0 | 1.246963 | 0.797 | 0.591 |
| CCL2 | 0 | 1.153067 | 0.201 | 0.041 |
| JUN | 0 | 1.143426 | 0.842 | 0.772 |
| C3 | 0 | 1.139472 | 0.84 | 0.456 |
| ICAM1 | 0 | 1.135248 | 0.651 | 0.483 |
| CXCL1 | 0 | 1.077755 | 0.293 | 0.097 |
| EGR1 | 0 | 1.072418 | 0.694 | 0.525 |
| SOD2 | 0 | 1.055837 | 0.769 | 0.603 |
| ERBB4 | 1.68E-110 | 1.041615 | 0.182 | 0.099 |
| JUNB | 0 | 1.02069 | 0.829 | 0.687 |
| RRAD | 0 | 0.946155 | 0.328 | 0.118 |
| DUSP1 | 0 | 0.927585 | 0.894 | 0.658 |
| ID1 | 0 | 0.912205 | 0.7 | 0.511 |
| SOX5 | 4.21E-265 | 0.909006 | 0.122 | 0.023 |
| MAPK10 | 0 | 0.870746 | 0.224 | 0.058 |
| KLF4 | 0 | 0.82082 | 0.326 | 0.084 |
| CD36 | 0 | 0.820471 | 0.504 | 0.246 |
| CXCL8 | 7.90E-165 | 0.775338 | 0.252 | 0.133 |
| CXCL3 | 1.91E-242 | 0.697865 | 0.284 | 0.133 |
| ID3 | 0 | 0.687593 | 0.393 | 0.198 |
| HSPA1A | 2.65E-168 | 0.639386 | 0.439 | 0.291 |
| ID2 | 7.55E-244 | 0.63303 | 0.465 | 0.305 |
| XBP1 | 0 | 0.626724 | 0.83 | 0.722 |
| TNFRSF1A | 0 | 0.584415 | 0.57 | 0.304 |
| UBC | 0 | 0.581075 | 0.906 | 0.877 |
| RAC1 | 0 | 0.577818 | 0.806 | 0.608 |
| HSPB1 | 0 | 0.558992 | 0.801 | 0.657 |
| PDZK1IP1 | 0 | 0.538204 | 0.481 | 0.235 |
| CDKN1A | 3.81E-273 | 0.531578 | 0.392 | 0.214 |
| HIF1A | 1.89E-295 | 0.494481 | 0.535 | 0.338 |
| PRKD1 | 4.29E-130 | 0.476476 | 0.125 | 0.049 |
| CEBPB | 0 | 0.450393 | 0.679 | 0.462 |
| TSC22D1 | 7.17E-226 | 0.445661 | 0.839 | 0.719 |
| BRAF | 2.85E-73 | 0.430835 | 0.272 | 0.189 |
| LMNA | 2.58E-113 | 0.426885 | 0.698 | 0.62 |
| IL32 | 0 | 0.418719 | 0.227 | 0.062 |
| SRSF2 | 1.94E-307 | 0.409407 | 0.65 | 0.469 |
| MIF | 2.57E-49 | 0.406996 | 0.63 | 0.627 |
| FGFR2 | 2.70E-25 | 0.393774 | 0.244 | 0.198 |

| | | | | |
|---|---|---|---|---|
| TOP1 | 1.03E-148 | 0.390084 | 0.796 | 0.704 |
| CD55 | 3.73E-48 | 0.371751 | 0.705 | 0.651 |
| PPP1R13B | 9.69E-103 | 0.367679 | 0.21 | 0.121 |
| SMC5 | 2.97E-120 | 0.356548 | 0.363 | 0.241 |
| ERN1 | 1.84E-150 | 0.343214 | 0.379 | 0.237 |
| NSMCE2 | 5.96E-45 | 0.336313 | 0.183 | 0.128 |
| FBP1 | 8.87E-233 | 0.336221 | 0.566 | 0.386 |
| KDM2A | 2.80E-92 | 0.330557 | 0.238 | 0.149 |
| SCMH1 | 1.61E-103 | 0.324639 | 0.137 | 0.065 |
| WWTR1 | 5.67E-23 | 0.318509 | 0.439 | 0.38 |
| DUOXA1 | 6.26E-17 | 0.314355 | 0.502 | 0.463 |
| SPRY4 | 6.71E-193 | 0.312054 | 0.37 | 0.214 |
| TEAD1 | 3.75E-97 | 0.311361 | 0.314 | 0.208 |
| DYRK1A | 6.33E-39 | 0.293529 | 0.225 | 0.168 |
| ZFP36L1 | 1.83E-32 | 0.292388 | 0.764 | 0.762 |
| ATXN1 | 0.0065199 | 0.286198 | 0.185 | 0.179 |
| IGFBP4 | 1.60E-182 | 0.272649 | 0.406 | 0.25 |
| BABAM2 | 1.12E-40 | 0.263499 | 0.224 | 0.166 |
| SMURF2 | 4.42E-55 | 0.260193 | 0.234 | 0.163 |
| PTTG1 | 1.97E-228 | 0.258005 | 0.211 | 0.082 |
| SOCS1 | 3.43E-297 | 0.257965 | 0.211 | 0.066 |
| CAT | 1.00E-169 | 0.257032 | 0.689 | 0.528 |
| SRSF3 | 7.21E-132 | 0.255689 | 0.74 | 0.628 |
| THRB | 2.24E-56 | 0.255262 | 0.146 | 0.089 |
| GPATCH8 | 7.30E-35 | 0.252401 | 0.195 | 0.144 |
| PHC2 | 2.52E-106 | 0.250933 | 0.312 | 0.205 |

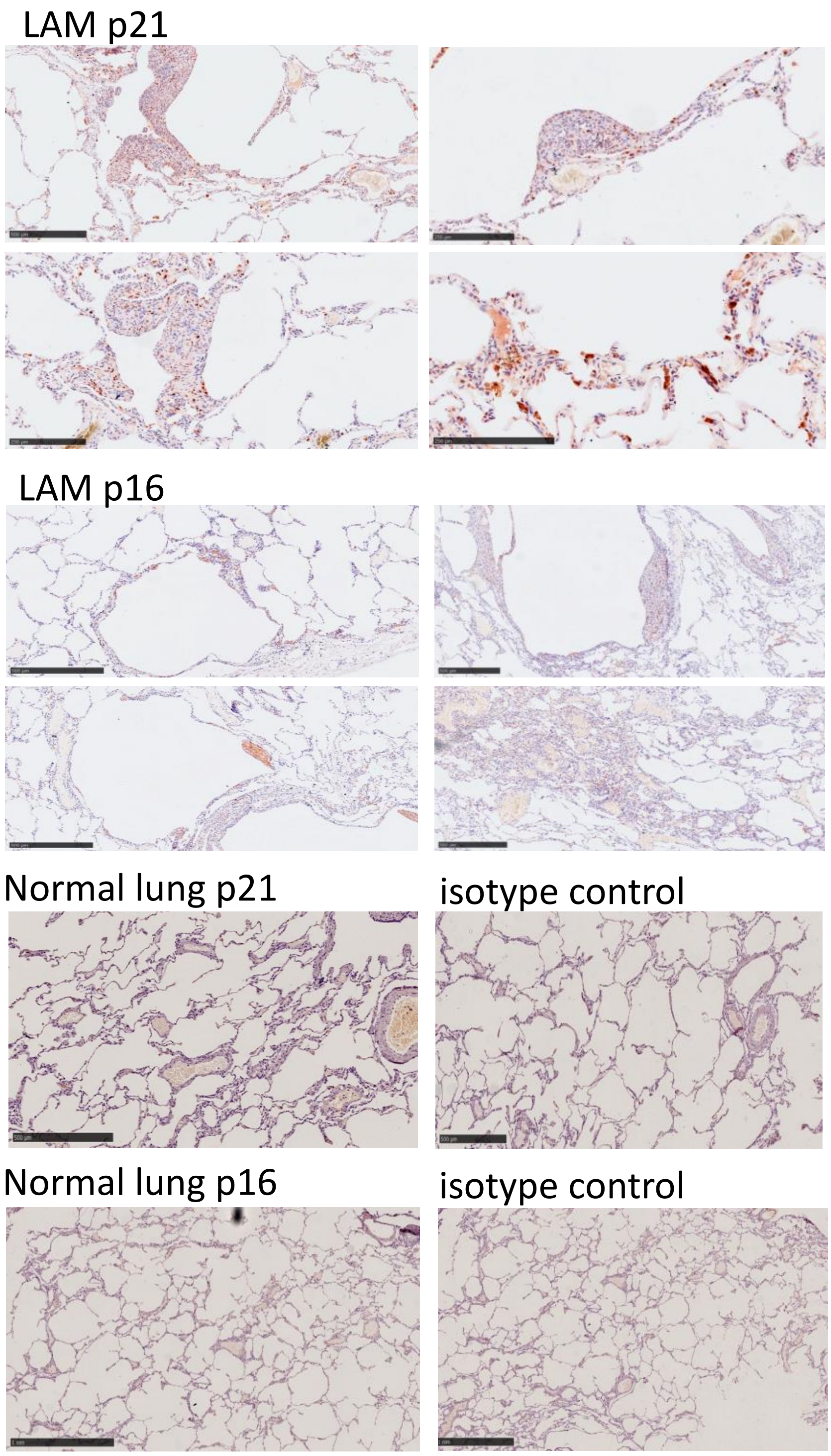


**Supplementary Figure 1. Immunohistochemical staining of LAM and control healthy lung tissue.** Representative images of senescence (p16 and p21), markers (brown) in LAM and healthy control lung tissue.

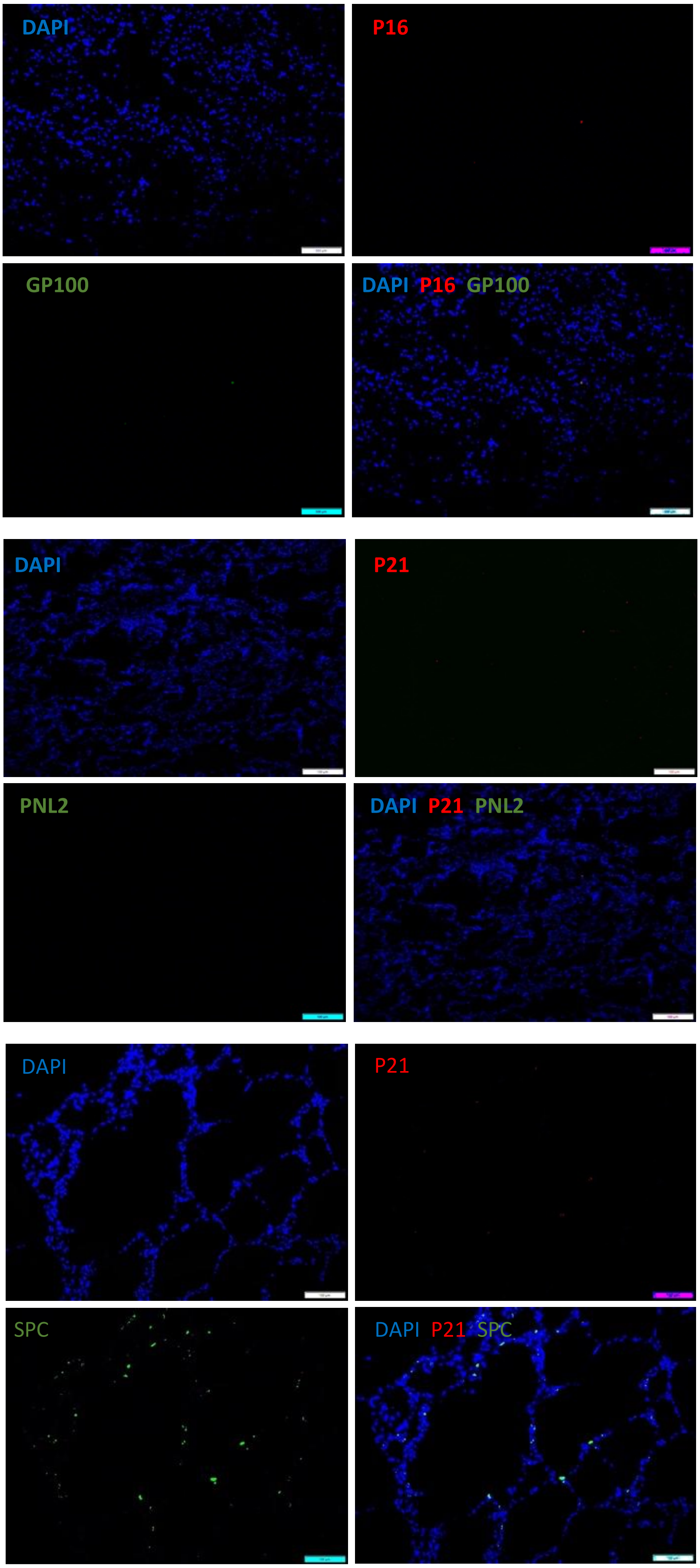


**Supplementary Figure 2. Immunohistochemical staining of control healthy lung tissue.** Representative images of senescence (p16 and p21), alveolar type 2 cell (SPC) and LAM cell (GP100 and PNL2) markers.

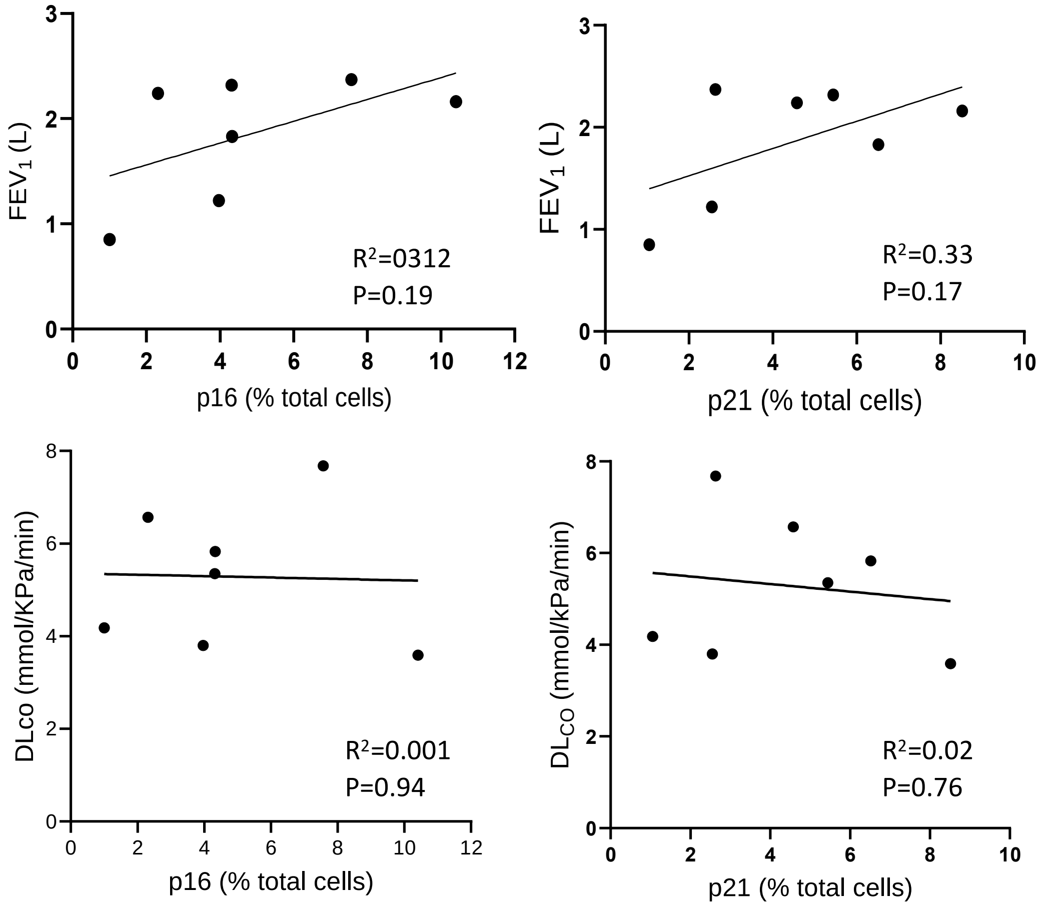


**Supplementary Figure 3. Relationship between histologic markers of senescence and lung function in women with LAM.** Percentage of p16 and p21 positive, compared with all cells in the region of interest in 7 LAM lungs correlated with forced expiratory volume in 1 second ($FEV_1$) and lung diffusion of carbon monoxide (DLco).

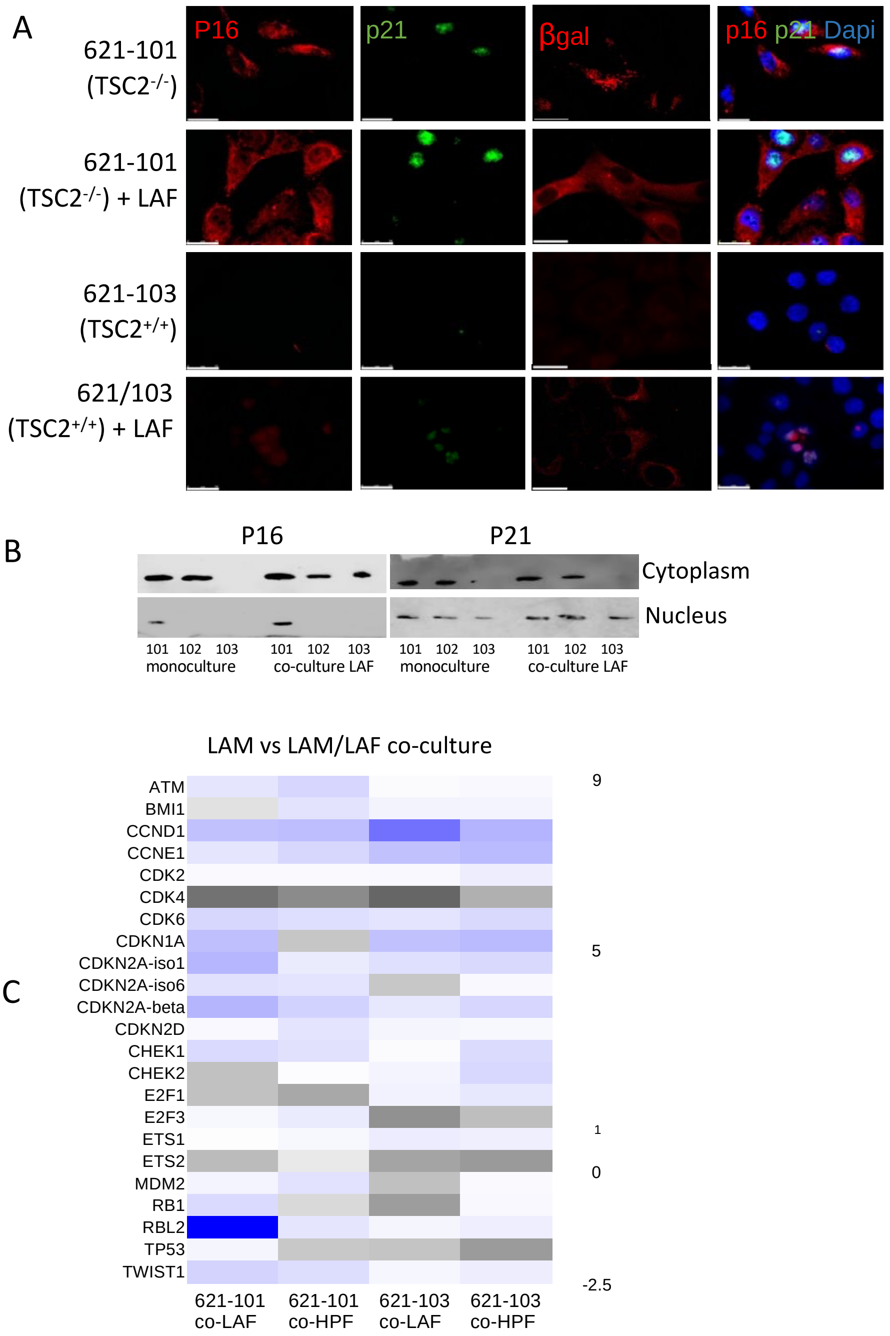


**Supplementary Figure 4. Markers of senescence in 621 cells and the effect TSC2.**
(A) Co-immunostaining of senescence markers p16, p21 and beta-galactosidase (βgal) in TSC2 $^{-/-}$ 621-101 and TSC2 $^{+/+}$ 621-103 cells alone or co-cultured with LAM associated fibroblasts (LAF) over 14 days. (B) Western blot of nuclear and cytoplasmic fractions of TSC2 $^{-/-}$ 621-101, TSC2 $^{+/+}$ 621-102 and TSC2 $^{+/+}$ 621-103 cells alone or co-cultured with LAFs probed for p16 and p21. (C) Heat map of senescence associated genes analysed by bulk RNA sequencing in TSC2 $^{-/-}$ 621-101 and TSC2 $^{+/+}$ 621-103 cells alone or co-cultured with either LAFs or normal human pulmonary fibroblasts (HPF).

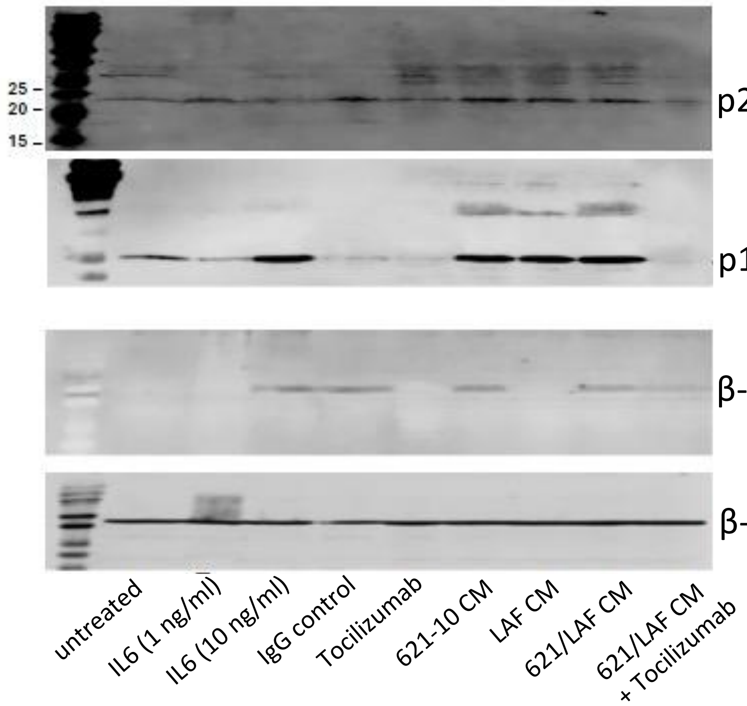


**Supplementary Figure 5. Markers of senescence in A549 cell/ LAM cell cocultures.** Western blot showing expression of p16, p21, and senescence associated β-galactosidase (SAβgal) protein expression by A549 cells over 14 days co-cultured with TSC2$^{-/-}$ 621-101 cells and LAM associated fibroblast (LAF) conditioned media (CM) in mono or co-cultures and the effect of Tocilizumab. β-actin is used as a loading control.

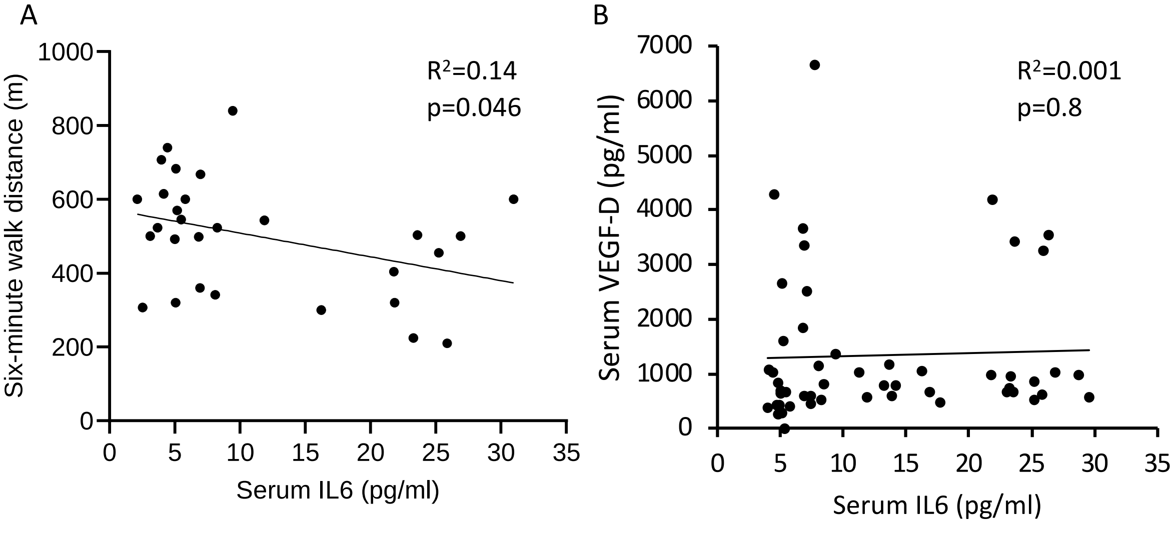


**Supplementary Figure 6. Association of IL6 with clinical features.**

A. Serum IL6 in rapamycin naive patients with LAM compared with six-minute walk distance. B. Serum IL6 in rapamycin naive patients with LAM compared with serum VEGF-D taken at the same time.
